# A two-dimensional space of linguistic representations shared across individuals

**DOI:** 10.1101/2025.05.21.655330

**Authors:** Greta Tuckute, Elizabeth J. Lee, Yongtian Ou, Evelina Fedorenko, Kendrick Kay

## Abstract

Our ability to extract meaning from linguistic inputs and package ideas into word sequences is supported by a network of left-hemisphere frontal and temporal brain areas. Despite extensive research, previous attempts to discover differences among these language areas have not revealed clear dissociations or spatial organization. All areas respond similarly during controlled linguistic experiments as well as during naturalistic language comprehension. To search for finer-grained organizational principles of language processing, we applied data-driven decomposition methods to ultra-high-field (7T) fMRI responses from eight participants listening to 200 linguistically diverse sentences. Using a cross-validation procedure that identifies shared structure across individuals, we find that two components successfully generalize across participants, together accounting for about 32% of the explainable variance in brain responses to sentences. The first component corresponds to processing difficulty, and the second—to meaning abstractness; we formally support this interpretation through targeted behavioral experiments and information-theoretic measures. Furthermore, we find that the two components are systematically organized within frontal and temporal language areas, with the meaning-abstractness component more prominent in the temporal regions. These findings reveal an interpretable, low-dimensional, spatially structured representational basis for language processing, and advance our understanding of linguistic representations at a detailed, fine-scale organizational level.

## Introduction

Language enables the transfer of complex ideas across minds—an ability that has laid a critical foundation for human culture. A set of left-lateralized frontal and temporal brain areas—the “language network”—supports language understanding and production, across modalities (spoken, written, and signed: MacSweeney et al. 2002; Menenti et al. 2011; Deniz et al. 2019; Giglio et al. 2022; Hu et al. 2022; Lipkin et al. 2022; Zada et al. 2024) and across typologically diverse languages (Malik-Moraleda et al. 2022). The language network responds in a highly selective manner to linguistic inputs (e.g., Fedorenko et al. 2011; Monti et al. 2012; Chen et al. 2023; see Fedorenko et al. 2024 for a recent review) and shows sensitivity to linguistic structure at different levels, from sound patterns to sentences (e.g., Vagharchakian et al. 2012; Regev et al. 2024; Shain, Kean et al. 2024a). The fact that these language areas support both comprehension and production suggests that they store our linguistic knowledge—including our knowledge of what words mean and how to put them together to create more complex meanings—which is needed for both interpreting and generating linguistic messages.

The internal organization of the language network remains poorly understood. What are the representational dimensions of this network, and do they exhibit a particular spatial organization? Many early proposals postulated distinctions at the regional level (e.g., Geschwind 1970; Dapretto & Bookheimer 1999; Hagoort 2005; Friederici 2012). However, a growing body of recent work has failed to find support for such dissociations (with the exception of the language area in the angular gyrus; e.g., Blank et al. 2016; Shain, Blank et al. 2020; Shain, Paunov, Chen et al. 2023). Instead, all language areas show similar response profiles, both in controlled experimental paradigms (e.g., Rodd et al. 2010; Fedorenko et al. 2020; Blank & Fedorenko, 2020; Hu et al. 2022) and during naturalistic comprehension (Shain, Blank et al. 2020; Wehbe et al. 2021; Shain et al. 2022). In particular, all language areas respond more strongly to structured and meaningful linguistic inputs compared to unstructured and meaningless ones (Fedorenko et al. 2010; Lipkin et al. 2022). For structured inputs, these areas are further modulated by linguistic complexity. For example, the level of BOLD responses during naturalistic comprehension can be predicted from incremental, word-by-word processing difficulty measures (Henderson et al. 2016; Wehbe et al. 2021) and from stimulus features, such as unexpected elements or unusual structures (e.g., Constable et al. 2004; Shain, Blank et al. 2020; Tuckute et al. 2024d) or non-local syntactic dependencies (Ben-Shachar et al. 2003; Blank et al. 2016; Shain et al. 2022).

However, some structure may exist within the language network that does not correspond to region-based distinctions. Indeed, functional differences among spatially interleaved neural populations have been reported within language areas in studies analyzing voxel-level fMRI responses or neuronal population activity recorded with invasive approaches. For example, using fMRI, Huth et al. (2016) reported differences in semantic tuning (preferences for linguistic input that express particular meanings) across the brain, including in frontal and temporal language areas (see also Deniz et al. 2019; Zhang et al. 2020). Using temporally-precise intracranial recordings, Regev, Casto et al. (2024) searched for differences in neural response profiles of the language network and found that neural populations differed in their temporal integration windows, varying between 1-2 words and 6-8 words (see also Jain et al. 2020 for related fMRI evidence). However, intracranial approaches only sample the cortex sparsely and from idiosyncratic locations across individuals, suggesting the need to systematically determine which representational dimensions are shared across individuals and how they map onto the broader cortical topography.

In this study, we set out to identify the organizing dimensions of language representations and to characterize how these dimensions are represented across the language network. Using ultra-high-field (7 Tesla) fMRI, we collected brain responses to 200 linguistically diverse sentences in eight proficient English speakers. We used a data-driven structure-discovery approach in which we explain each voxel’s response to diverse sentences as a weighted sum of a small number of components (see e.g., Norman-Haignere et al. 2015; Khosla et al. 2022, for similar approaches in other domains), and then systematically tested which components of language representations generalize across individuals. To foreshadow the critical results, we found a two-dimensional space that is shared across individuals and explains substantial variance in voxel-level responses: one axis corresponds to processing difficulty, and the other—to meaning abstractness. Surprisingly, we also found that the two components could account for activity in parts of the visual ventral stream, traditionally not considered part of the language network (cf. Li et al. 2024; Wolna et al. 2025). The two components furthermore exhibited systematic topography across participants, revealing spatially fine-grained organization of language processing.

## Results

Our results are organized according to three questions: (1) How many distinct components of language representations are shared across individuals? (2) What characterizes the components? and (3) How are the components spatially organized?

### 1. How many components of language are shared across individuals?

We collected ultra-high-field (7 Tesla) fMRI responses to a set of 200 linguistically diverse sentences from eight participants (**Figure 1A-B**). All participants were native or proficient English speakers; six participants were also proficient in one or two languages other than English (<u>Methods; Participants</u>). To search for the main organizing dimensions of language representations, we took a data-driven approach and performed singular value decomposition (SVD) of brain responses to obtain principal components of variance (**Figure 1C-E**). The SVD was applied to the portion of data covariance that is attributable to sentence-evoked responses (signal), ignoring trial-to-trial variability (noise) (<u>Methods; Deriving Sentence PCs)</u>. We denote the derived components as ‘Sentence PCs’, where each Sentence PC indicates how much each sentence drives variance along a given direction in voxel space.

**Figure 1.**
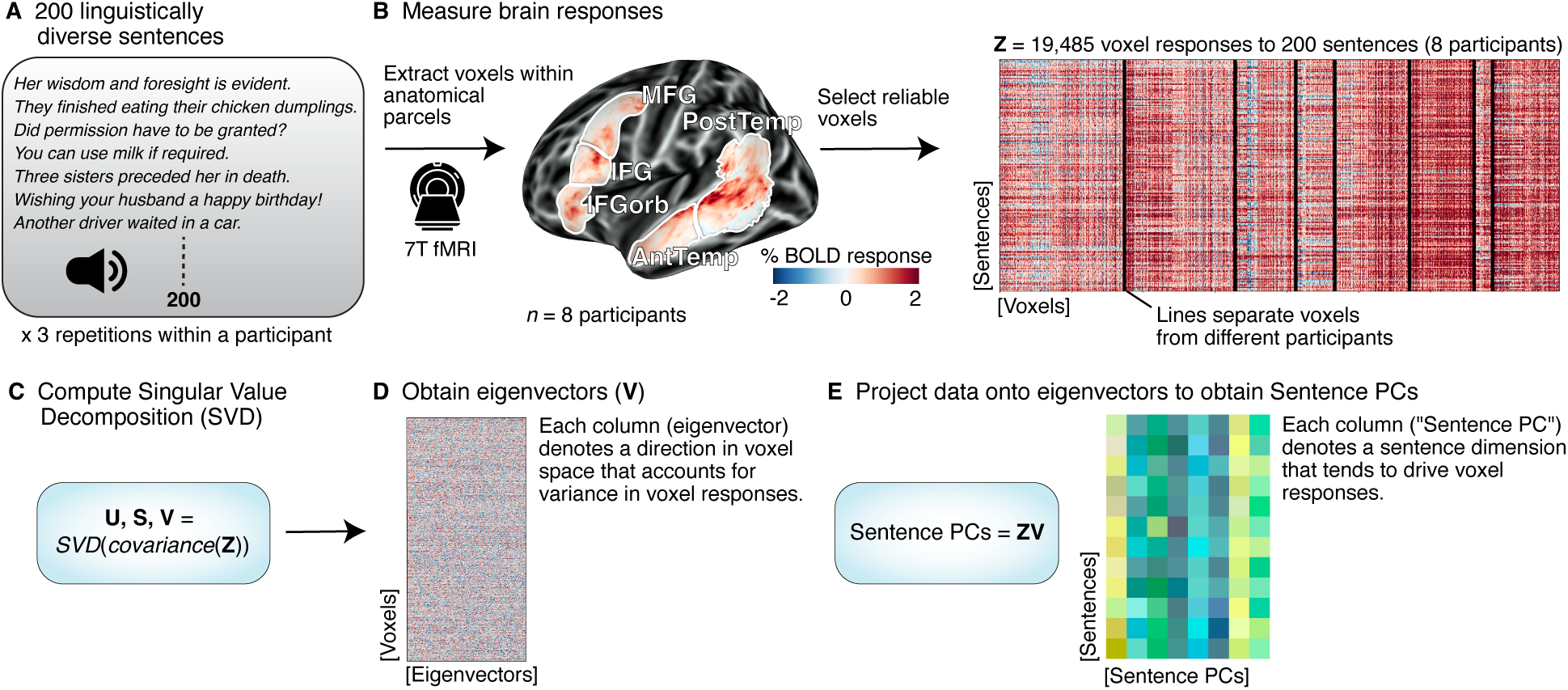
Procedure for deriving Sentence Principal Components (PCs). (**A**) Eight participants listened to 200 linguistically diverse sentences during fMRI scanning, with each sentence repeated three times in a pseudorandomized order. **(B)** Brain responses to each sentence were extracted within five broad anatomical parcels (‘language parcels’) within which most or all individuals in prior studies showed responses to language in a validated language localizer contrast, including in large samples (e.g., Fedorenko et al. 2010; Lipkin et al. 2022; see <u>Methods; Language parcels</u>). Voxels were aggregated across eight participants, resulting in a data matrix **Z**. In the primary analyses, we included voxels that showed reliable responses to the experiment (voxels with a noise ceiling signal-to-noise ratio above 0.4; <u>Methods; Deriving Sentence PCs</u>) (see <u>SI 1</u> for evidence that the results are robust to different voxel inclusion criteria). (**C**) We applied Generative Modeling of Signal and Noise (GSN; Kay et al. 2024) to isolate the covariance of the signal in the data matrix, excluding noise defined as variability across trial repetitions. We then applied singular value decomposition (SVD) to the signal covariance (see SI 1 for analyses that use the standard covariance matrix, which are quantitatively similar). (**D**) The SVD yielded eigenvectors (V in the schematic) which represent directions in voxel space along which the data exhibits most variance. (**E**) The data matrix, Z, was projected onto the eigenvectors, V, to obtain the sentence projections (denoted as ‘Sentence PCs’). Each Sentence PC denotes how much each sentence drives variance along a principal dimension of voxel activity (across language parcels and participants).

Because we are interested in identifying the components that are *shared* across individuals, we performed a nested cross-validation to estimate the number of generalizable Sentence PCs (**Figure 2A**). In each iteration, we derived Sentence PCs using data from all participants except one (i.e., seven participants) and evaluated the generalizability of the estimated Sentence PCs by predicting single-voxel responses of the held-out participant using ordinary least squares (OLS) regression. Each voxel’s response was modeled as a weighted sum of Sentence PCs and evaluated through the coefficient of determination (*R*^2^)—a stringent measure of how well the predicted responses match the measured voxel responses in terms of both their mean and variance. **Figure 2B** shows the *R*^2^ performance across participants in the broad anatomical language parcels. Model performance peaks with two Sentence PCs (mean *R*^2^ = 11.91% +/- SEM across participants 0.90%), corresponding to 28.94% of the noise ceiling—the theoretical upper limit for model performance (average noise ceiling = 41.16%; horizontal gray lines in **Figure 2B**). Including additional components beyond two does not further improve prediction accuracy, indicating that additional components beyond two do not generalize to new participants. The two Sentence PC result was highly robust to different criteria for estimating the Sentence PCs (<u>SI 1</u>) and different voxel inclusion criteria for evaluation (<u>SI 2</u>).

**Figure 2.**
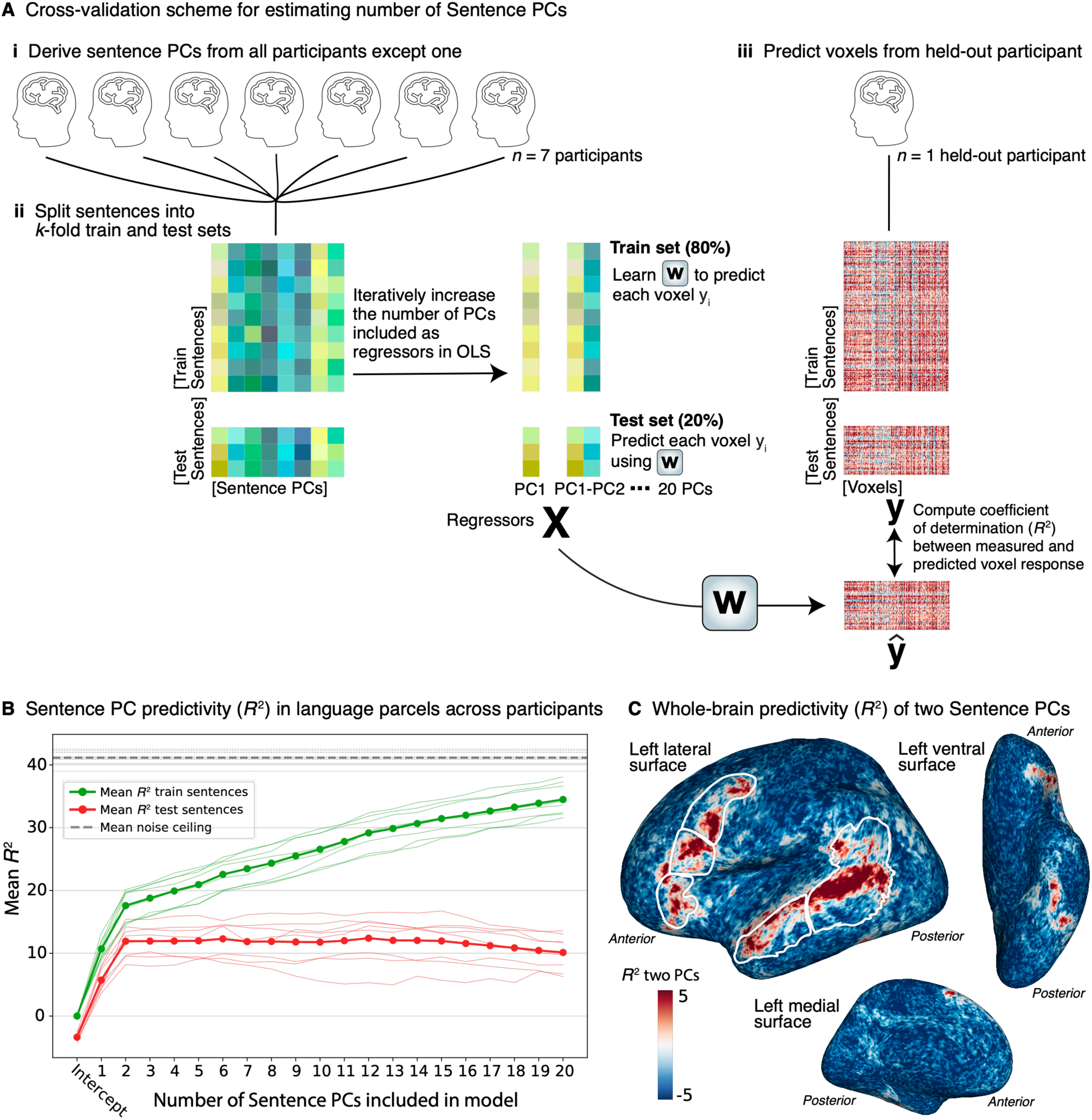
Number of components (‘Sentence PCs’) that generalize across individuals. (**A**) To assess how many Sentence PCs generalize across participants, we estimated Sentence PCs from all participants except one (n = 7) and used these PCs to predict voxel responses from the held-out participant. For each held-out participant, OLS regression was performed by iteratively increasing the number of Sentence PCs included as regressors, X. This procedure was carried out in a k-fold cross-validation scheme (k = 5, results remained quantitatively similar for k = 2 or k = 8), where a subset of sentences was used to estimate the weights (w), which were then applied to predict responses for an independent set of sentences. In each cross-validation split, we computed the coefficient of determination (R2) between the predicted and measured voxel responses (see more details in Methods; Modeling voxels using Sentence PCs). **(B)** Predictivity (coefficient of determination, *R*^2^) for models with an increasing number of Sentence PCs used as regressors to predict voxels in a held-out participant. Thin colored lines show *R*^2^ for individual participants; thick lines show the mean. Red lines show the *R*^2^ test performance, green lines show the *R*^2^ train performance. The analysis included evaluation on voxels extracted from the broad anatomical language parcels (see outlines in panel **C**) that passed a reliability threshold of a noise ceiling signal-to-noise ratio of 0.4 (an average of 2,368 voxels per participant +/- SEM 1,173). The horizontal gray lines show the noise ceiling: the thick line shows the average noise ceiling across participants, thin lines show noise ceilings for individual participants. See <u>SI 2</u> for evaluation on voxels with different or no reliability inclusion criteria; critically, the finding that two components are optimal remains consistent regardless of voxel inclusion criteria. **(C)** Predictivity (*R*^2^), averaged across *n* = 8 participants, plotted on the inflated left hemisphere surface (in the fsaverage space; Fischl et al. 1999). The white outlines on the surface map delineate the anatomical language parcels (<u>Methods; Language parcels</u>).

How much variance in brain responses to sentences do the two identified components account for? Our two analyses provide a converging answer. From the SVD analysis, by examining the cumulative sizes of the eigenvalues, we find that the two components account for 34.05% of the signal covariance. From the OLS analysis, we find that the two components predict 28.94% of the noise ceiling in the held-out participants. Both analyses discount measurement noise and reflect repeatable, sentence-evoked signals. Thus, the two Sentence PCs define a robust two-dimensional space shared across participants that accounts for about 32% of sentence-evoked brain responses.

The quantifications described in **Figure 2B** target the anatomical language parcels. For a better sense of the spatial specificity of the two Sentence PC model, we plot a whole-brain surface map showing *R*^2^ performance in the left hemisphere, averaged across participants (**Figure 2C**). Performance is highest in frontal and temporal areas traditionally associated with language processing (e.g., Binder et al. 1997; Lipkin et al. 2022; see also <u>SI 3</u> for correlation of model predictivity with language selectivity). In addition to the lateral frontal and temporal areas, we observe significant prediction performance in the left ventral temporal cortex (VTC) (<u>SI 3</u>). The two Sentence PCs could also predict responses in the right hemisphere frontal and temporal areas, albeit performance was weaker than in the left hemisphere, and largely absent in the right VTC (<u>SI 4</u>).

### 2. What characterizes the two sentence components?

Having identified two components that robustly emerge from the sentence-evoked brain responses and are shared across individuals, we next sought to interpret these components to understand the main dimensions of language representations in the brain. We estimated Sentence PCs using data from all eight participants, visualized the 200 sentences in the two-dimensional space spanned by Sentence PC 1 and PC 2 (**Figure 3A**), and examined their correlation with linguistic/semantic properties (**Figure 3B**).

**Figure 3.**
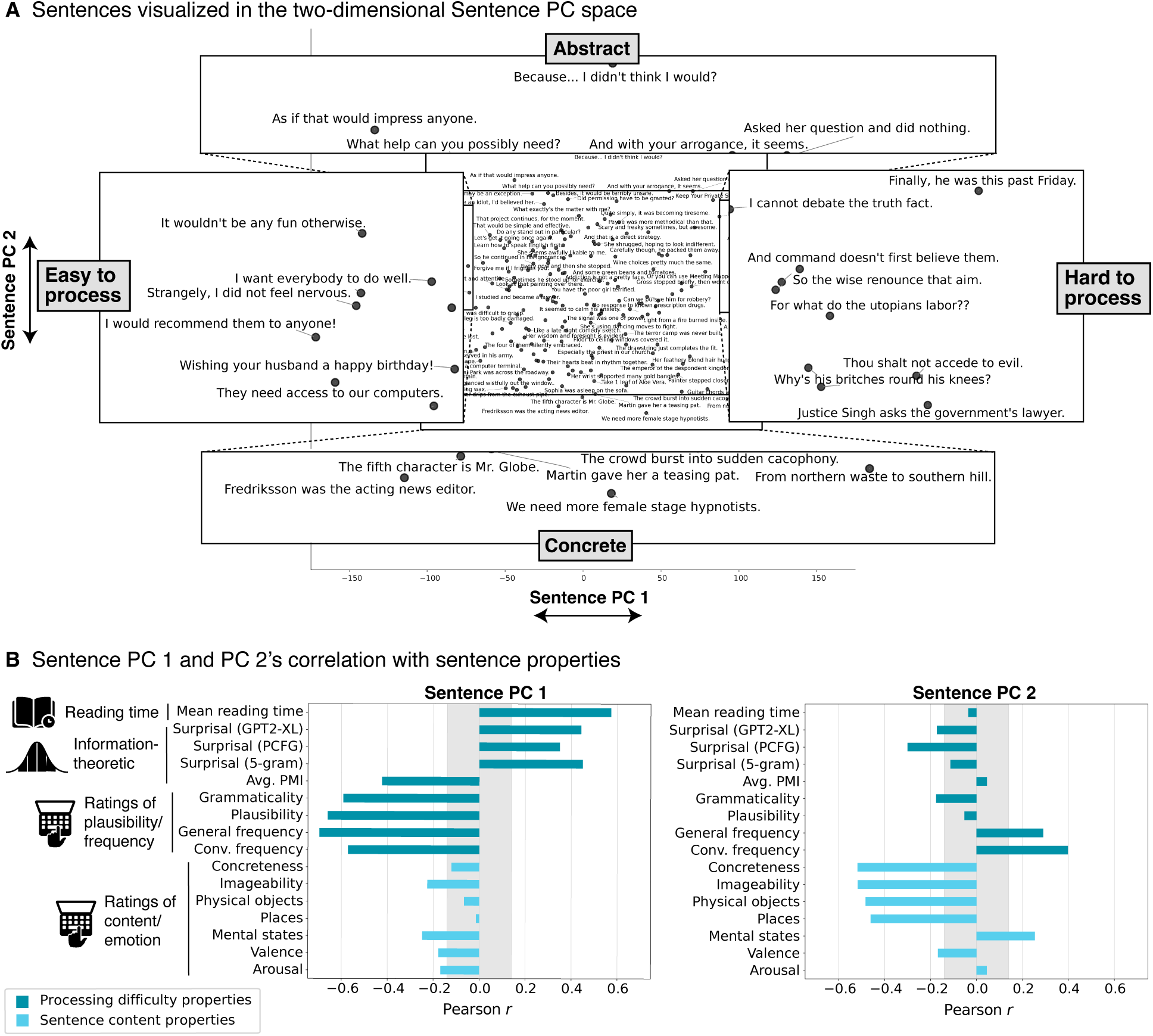
Visualization and characterization of Sentence PC 1 and PC 2. (**A**) Individual sentences (with a few sentences omitted for visibility) plotted in the two-dimensional space spanned by Sentence PC 1 (x-axis) and Sentence PC 2 (y-axis). The insets zoom in on the lowest and highest values observed for each Sentence PC. (**B**) Correlation of Sentence PC 1 and PC 2 with 16 linguistic/semantic properties within two broad categories—linguistic processing difficulty (teal bars) and sentence content (light blue bars)—measured through behavioral experiments and information-theoretic estimates. The vertical gray bar marks range of correlation values expected by chance for a two-tailed p<.05 threshold based on 200 samples.

A visualization of the two-dimensional space is shown in **Figure 3A**. It appears that along PC 1, sentences differ in well-formedness, while along PC 2, they vary in content. To quantitatively evaluate intuitions about these dimensions, we correlated the two Sentence PCs with 16 sentence properties, all motivated by past work on language processing (<u>Methods; Sentence properties</u>). The properties can be grouped into two broad categories: properties related to processing difficulty (nine properties) and properties related to sentence meaning (seven properties). The first group of properties includes behavioral and computational measures that reflect how effortful a sentence is to process. The first property, average reading time, was measured using a Maze reading task (Forster et al. 2009; Boyce et al. 2020), where participants (independent from those in the fMRI study) chose, at each position in a sentence, between a word that fits the preceding context (the actual continuation of the sentence) and a distractor that does not. Response times provide an online measure of processing effort (Forster et al. 2009; Boyce & Levy 2023; Hao et al. 2023), with slower times corresponding to parts of the sentence that are more difficult to understand. We averaged word-level reading times across each sentence to obtain a sentence-level measure (see <u>Methods; Word-by-word reading times</u> for experimental details). The next three properties in this group were motivated by a large body of work demonstrating the importance of surprisal—the degree of contextual predictability—in explaining human behavior and neural responses during language processing (Demberg & Keller 2008; Smith & Levy 2013; Henderson et al. 2016; Willems et al. 2016; Brothers & Kuperberg 2021; Heilbron et al. 2022; Shain, Blank et al. 2020, Shain et al. 2024b). We estimated surprisal using three models: a GPT2-XL language model trained on large text corpora (Radford et al. 2018), a lexicalized probabilistic context-free grammar (PCFG) model that focuses on the syntactic structure of sentences (van Schijndel et al. 2013), and an *n*-gram model, which estimates predictability based on the preceding four words. We also included pointwise mutual information (PMI) to quantify the association among nearby words based on co-occurrence counts (Church & Hanks 1990), which has been shown to modulate responses in the language network (Mollica, Siegelman et al. 2020). Finally, we obtained behavioral ratings of each sentence’s grammatical well-formedness (How much does the sentence obey the rules of English grammar?), semantic plausibility (How much sense does the sentence make?), and perceived frequency of occurrence in general and in conversational contexts. As expected, many of these properties are somewhat correlated with each other. For example, as also demonstrated in prior work (Rayner et al. 2006; Smith & Levy 2013; Shain et al. 2024b), we found that infrequent and surprising sentences tend to take longer to read (<u>SI 5</u>).

The properties in the second group all capture aspects of a sentence’s meaning. These properties were motivated by prior work (e.g., Paivio et al. 1968; Kuchinke et al. 2005; Binder et al. 2005; Jack et al. 2013; Huth et al. 2016; Arfé et al. 2022), including the semantic tuning effects mentioned in the introduction (Huth et al. 2016), and general preferences of the language areas for more abstract content (Noppeney & Price 2004; Tuckute et al. 2024d). All properties in this group were derived from behavioral ratings. We included ratings of concreteness (How much is a sentence’s meaning tied to perceptual experience?) and imageability (How easy is a sentence to visualize?). Three properties targeted the content of each sentence: How much does the sentence make you think about (1) physical objects and their interactions, (2) places and environments, and (3) others’ mental states? Finally, we assessed two emotional dimensions of each sentence: valence (How positive is the sentence’s content?) and arousal (How exciting is the sentence’s content?) (see Tuckute et al. (2024d) and <u>Methods; Behavioral norms</u>). As expected, concreteness and imageability were highly correlated (Pearson *r* = 0.85, *p*<.001), and both were also positively correlated with whether a sentence’s content is about physical objects and places (*r*s = 0.47–0.79, *p*s<.001), but not mental states (*r*s = 0.03–0.06, n.s.). In contrast, the emotional dimensions— valence and arousal—were only weakly correlated with these other content-based ratings (*r*s = - 0.03–0.12, n.s.), except that sentences with mental state content were positively correlated with arousal (*r* = 0.53, *p*<.001) (<u>SI 5</u>).

Figure 3B shows that Sentence PC 1 is strongly correlated with the linguistic properties related to processing difficulty (teal bars), including reading times, surprisal, and behavioral ratings of linguistic well-formedness and frequency. The numerically strongest correlation is with the “General frequency” measure (*r* = −0.70, *p*<.001), a measure of people’s perception of how likely they are to encounter a given sentence. PC 1 is also strongly correlated with average reading time (*r* = 0.57, *p*<.001), meaning that sentences with high loadings on this PC take longer to process, establishing a link between neural response tuning and behavioral online processing effort (see also Wehbe et al. 2021). We label this dimension “Processing difficulty”: sentences with high loadings on this PC require more effort to read and understand, are more surprising, and are perceived as less frequent.

For Sentence PC 2, the strongest correlations are with properties related to whether a sentence’s content is about physical objects/places, and whether a sentence is tied to perceptual, including visual, experience. The numerically strongest correlation is with the “Concreteness” measure (*r* = −0.52, *p*<.001)—how much a sentence is tied to perceptual experiences (vision, audition, smell, touch, etc.) (Reilly et al. 2025). We therefore label this dimension “Abstractness”: sentences with high loadings on this PC are hard to visualize and convey abstract content.

In summary, the two Sentence PCs—derived solely from the decomposition of brain responses to a set of linguistically diverse sentences—are highly interpretable, yielding insights into the core dimensions of language processing.

### 3. How are the components spatially organized?

Having established which linguistic and semantic properties characterize the two Sentence PCs, we next investigated their spatial organization in the brain to understand how these components are distributed and whether any systematic patterns exist across individuals. Specifically, we analyzed the voxel-wise weights learned for the two Sentence PCs during OLS regression (Figure 2A and <u>Methods; Modeling voxels using Sentence PCs</u>). These weights place each voxel within a two-dimensional space that specifies the contributions of the two Sentence PCs in accounting for the voxel’s responses to the sentences. In this space, the *direction* (angle) of a voxel reflects the relative influence of the two components in accounting for the voxel’s response, whereas the *magnitude* along that direction reflects the voxel’s response units (BOLD percent signal change). We focus on the voxel-level angles as it allows us to assess the contributions of the two Sentence PCs independently of general fluctuations in BOLD magnitude (affected by many factors, including proximity to blood vessels). These voxel-level angles showed high reliability as estimated via split-half analyses across independent sets of stimuli (<u>SI 7</u>).

To aid visualization, we divided the two-dimensional sentence space into 16 sectors and assigned colors to each sector using a “rainbow” color scheme (Figure 4A). Along Sentence PC 1 (x-axis), green colors denote “Hard to process” sentences while purple colors denote “Easy to process” sentences. Along Sentence PC 2, blue colors denote the “Abstract” sentences, while red colors denote “Concrete” sentences. The surface maps in Figure 4A use this 16-sector color scheme to visualize the average angle across *n* = 8 participants. For the group-average map, a voxel is included if at least three participants show significantly predicted activity at that voxel location (Figure 4E shows examples of individual participant maps; see <u>SI 8</u> for maps for the remaining participants). The surface maps reveal that the Sentence PCs are systematically distributed across the lateral and ventral surface, forming a large-scale topography (see also <u>SI 9</u> for consistency of the surface maps and angle sectors across independent splits of participants).

**Figure 4.**
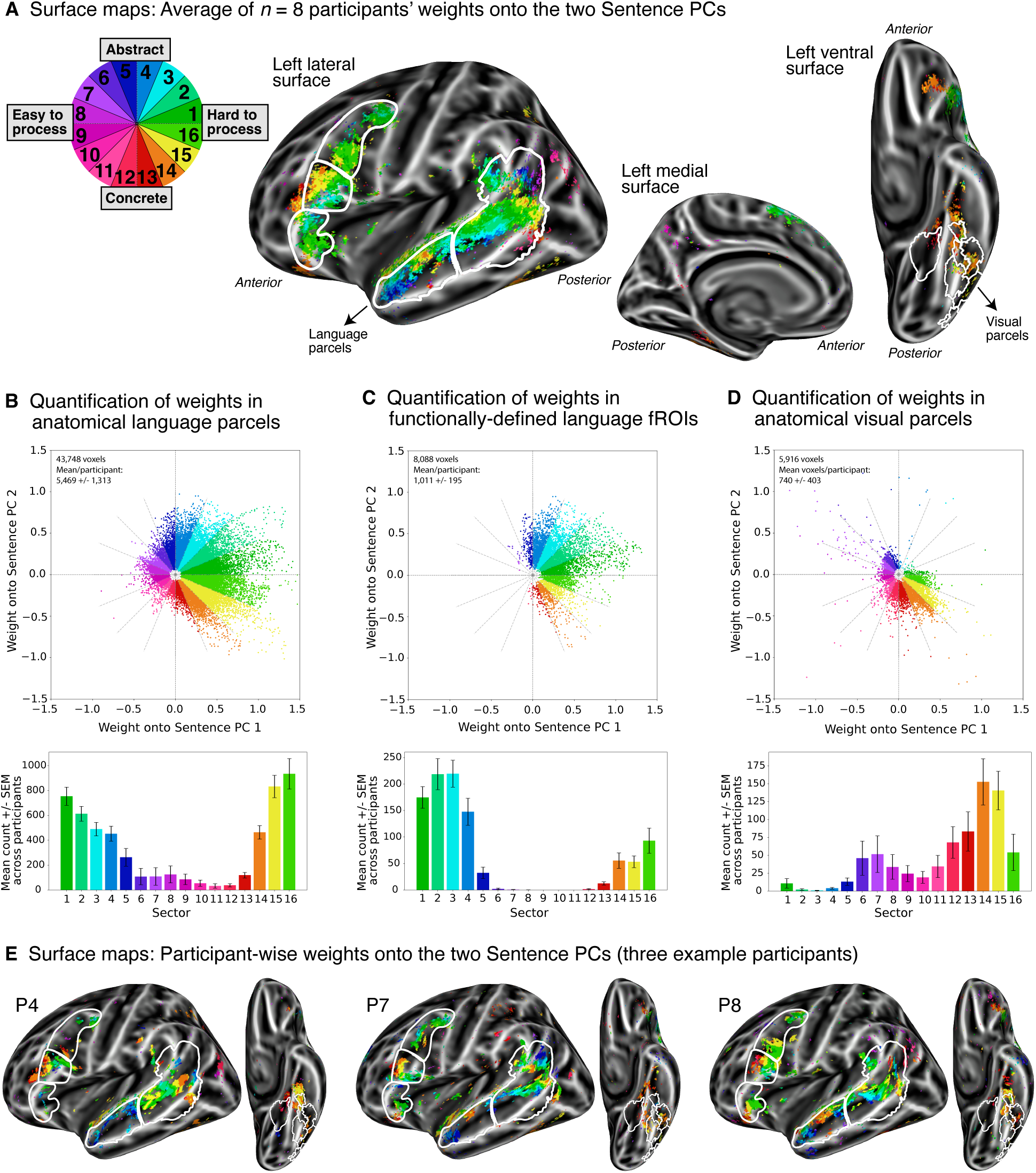
Whole-brain spatial organization of the two Sentence PC-model. **(A)** The surface maps show the average of *n* = 8 participants’ weights onto the two Sentence PCs for significantly predicted voxels. In the OLS analysis, each voxel gets weighted along the two PCs, indicating the contribution of each PC to its responses. To interpret the relative contribution of the PCs, we use a “rainbow” colormap where the weights for a given voxel define an angle (direction) which we color-code according to 16 distinct sectors. In the group-averaged surface map, voxels were included if at least three participants exhibited a significantly predicted voxel (<u>Methods; Modeling voxels using Sentence PCs</u>). **(B-D)** Quantification of the angle sectors for voxels within five anatomical probabilistic language parcels (Lipkin et al. 2022) (panel **B**), five functionally-defined language fROIs (Fedorenko et al. 2010) (panel **C**), and ten anatomical probabilistic visual parcels (Rosenke et al. 2021) (panel **D**). The scatter plots show the weights for each voxel onto Sentence PC 1 (x-axis) and Sentence PC 2 (y-axis) across *n* = 8 participants for significantly predicted voxels (in panel **B**, 12 voxels are not shown on the scatter plot as they fell outside the axis limits, and in panel **D**, 2 voxels not shown). The histograms show the average number of voxels within each of the 16 sectors in the two-PC space (using the color scheme illustrated in panel **A**). Error bars denote SEM across participants. (**E**) Surface maps for three individual participants (see SI 8 for all participants). As in panel A, the maps show the weights onto the two Sentence PCs using the rainbow sector color scheme. Voxels that were significantly predicted are shown.

To quantify the Sentence PCs’ spatial organization and to understand their distribution in well-studied frontal and temporal language areas (Binder et al. 1997; Fedorenko et al. 2010; Lipkin et al. 2022), we plot histograms of voxel-level angles in three sets of voxels: those in (1) five anatomical language parcels, derived from a probabilistic map of language-selective areas across 220 independent individuals (Figure 4B; <u>Methods; Language parcels</u>), and (2) five functionally-defined language fROIs representing the most language-selective voxels within each participant (Figure 4C; <u>Methods; Language network fROIs</u>). Additionally, motivated by our finding that the two Sentence PCs could also predict activity in the left VTC (Figure 2C), we investigated these language-evoked responses in voxels across (3) ten anatomical probabilistic parcels corresponding to visual category-selective regions (Figure 4D; <u>Methods; Visual parcels</u>) (Rosenke et al. 2021). The scatter plots in Figure 4B-D illustrate each voxel’s position in the two-dimensional sentence space within each of the three sets of voxels, and the histograms display voxel count within each angle sector, aggregated across participants.

First, considering the five language parcels (three in the frontal lobe: orbital part of IFG, IFG, MFG; and two in the temporal lobe: anterior and posterior), we see a clear preference for hard-to-process/abstract sentences (Figure 4B and Figure 5A). The anterior temporal parcel contains a large proportion of voxels with preference for abstract sentences (blue colors). Statistically, it is significantly different from the IFG and MFG parcels (both *p*<.05, determined via a two-tailed non-parametric test which permutes the parcel assignment of the participant-level circular averages) and shows a trend toward significance from the IFG orbital parcel (*p*=0.058). The posterior temporal parcel is significantly different from IFG (*p*<.05) and shows a trend for being different from MFG (*p*=0.058). The frontal parcels do not differ significantly from one another, nor do the two temporal parcels differ from each other.

**Figure 5.**
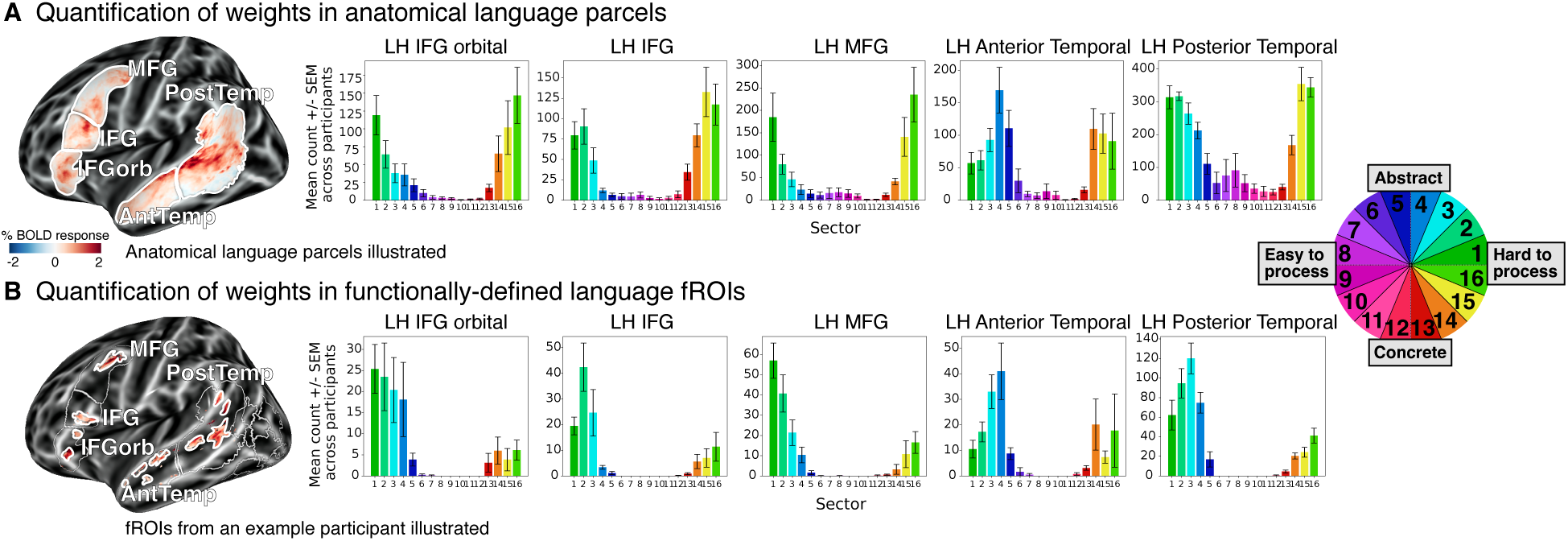
Quantification of Sentence PC weights in each language parcel and each language fROI. Quantification of angle sectors for voxels within each of five anatomical probabilistic language parcels (Lipkin et al. 2022) (panel **A**; the count across all five parcels is shown in **Figure 4B**) and within each of five functionally-defined language fROIs (Fedorenko et al. 2010) (panel **B**; the count across all five fROIs is shown in **Figure 4C**). Error bars denote SEM across participants. The surface map in panel **A** illustrates the anatomical language parcels, and the surface map in panel **B** illustrates the fROIs from an example participant (the precise fROI locations differ across participants). The fROIs were defined as the top 10% most significant voxels for a *sentences > nonwords* contrast, identified using a well-validated localizer task (e.g., Fedorenko et al. 2010; Lipkin et al. 2022) within the anatomical parcels analyzed in panel **A** (<u>Methods; Language network fROIs</u>).

Next, we zoomed in on the functionally-defined language reions in each individual (functional regions of interest (fROIs)) (Figure 4C and Figure 5B). These fROIs were defined by taking the top 10% most language-responsive voxels based on a well-validated language localizer (using a *sentences* > *nonwords* contrast; Fedorenko et al. 2010) in the aforementioned anatomical parcels. These fROIs have been extensively studied (e.g., Fedorenko et al. 2012; Ivanova et al. 2020; Braga et al. 2020; Wehbe et al. 2021; see Fedorenko et al. 2024 for a review). Strikingly, the fROIs all fall strictly in the hard-to-process/abstract part of the space, with almost no voxels in other parts of the space (see also <u>SI 11</u>). None of the fROIs differ significantly from one another, with the one exception of the posterior temporal fROI being different from the IFG and MFG fROIs (*p*s<.05).

How can we relate these findings of similar fROI tuning to the statistical differences observed among anatomical language parcels? The latter differences are plausibly due to the inclusion of voxels from nearby functional networks known to be proximal to the language network (Braga et al. 2020; Fedorenko & Blank 2020; DiNicola et al. 2023). Thus, the language-selective regions carve out a particular part of the tuning landscape, whereas surrounding cortical areas—included within the broader anatomically-defined language parcels—have more diverse tuning properties (Figure 4B and Figure 5A).

Finally, we examined the significantly predicted voxels in the VTC (Figure 2C). This part of the brain is traditionally associated with high-level visual processing, including category-selective areas involved in recognizing faces, bodies, and places (Kanwisher et al. 1997; Epstein & Kanwisher 1998; Peelen & Downing 2005; Grill-Spector & Weiner 2014). In an attempt to further characterize the specific sub-regions within this area, we leveraged a probabilistic atlas for category-selective visual regions, such as FFA-1, FFA-2, EBA, and VWFA-1 (Rosenke et al. 2021; <u>Methods; Visual parcels</u>). Figure 4D shows the angle sector counts of significantly predicted voxels within these visual parcels. These regions exhibit a markedly different profile compared to both the language parcels and fROIs (*p*s<.005), with the visual parcels showing a preference for concrete sentences (see also <u>SI 11</u>). In a supplemental analysis, motivated by prior findings of language-selective regions in the VTC (Lüders et al. 1991; Li et al. 2024; Wolna et al. 2025), we examined the angle sectors within a probabilistic language parcel in the VTC derived from a previous study (<u>Methods; Ventral temporal language parcel</u>; Li et al. 2024) (partly overlapping with category-selective parcels). In line with the findings above, we find that the responses in this area are similarly tuned to concrete sentences, but the well-predicted voxels extend beyond the probabilistic language parcel (<u>SI 10</u>). These results show that the two-dimensional organization of linguistic representations extends beyond the traditional fronto-temporal language network into the left-hemisphere VTC (but not right-hemisphere VTC; <u>SI 4</u>), revealing a specific tuning for concrete, imageable sentences during language comprehension.

## Discussion

The novel contributions of this study are threefold. First, through data-driven decomposition of brain responses to linguistically diverse sentences, we showed that only two components—shared across individuals—emerged robustly, together accounting for about 32% of the variance driven by sentences. Analysis of the sentences’ linguistic and semantic properties revealed that the first component corresponds to processing difficulty, and the second—to meaning abstractness. Second, we found that all frontal and temporal language areas respond more strongly to hard-to-process sentences that express abstract meanings, with the temporal areas being more tuned to abstract meanings compared to the frontal ones. And third, we demonstrated that these two components exhibit systematic topographic organization across frontal and temporal brain areas, and extend into the left ventral visual stream. These results provide a spatially precise view of the shared dimensions underlying language comprehension in the human brain. Below, we elaborate on these findings.

### 1. Are two components all you need?

Although every child growing up in a linguistic environment acquires one or more languages, human linguistic experiences differ substantially, including in the amount and kind of linguistic input received across the lifespan (Gilkerson et al. 2017; Dailey & Bergelson 2022; Bergelson et al. 2023). In spite of this variability, speakers of the same language can communicate effectively about almost any topic. What are the common neural dimensions that support language processing across minds? In this study, we found that only two components—processing difficulty and meaning abstractness—are common to linguistic representations across individuals. Of course, these findings do not imply that these dimensions are sufficient for language comprehension. For example, the sentences “My skin feels like melting wax.” and “Water drips from the exhaust pipe.” occupy a similar position in the two-dimensional space of processing difficulty and meaning abstractness, yet clearly carry distinct meanings.

A priori, two components might seem like surprisingly few. Our analyses show that these components account for a substantial amount of variance, but some variance is left unexplained. Our main approach—singular value decomposition of sentence-evoked voxel responses across participants—revealed that the two components explain 34% of the variance driven by sentences. A complementary approach—leave-one-participant-out regression—produced a similar estimate: the two components predict 29% of the noise ceiling (i.e., the maximum amount of variance that can in theory be predicted, given measurement noise). From these analyses, we can conclude that about a third of the variance driven by sentences reflects shared linguistic representations and that the remaining two thirds (66-71%) pertain to inter-individual variability that does not generalize across individuals. Future work could examine the nature of these participant-unique representations: perhaps they reflect semantic representations that are shaped by individual experience (see also <u>Discussion; 4. What are the two Sentence PCs representing and computing?</u>).

The data-driven analysis approach we implemented removes the unwanted effects of measurement noise and identifies the highest-variance components that are shared across individuals. Nonetheless, there are experiment-related factors that may have obscured potential additional shared components. First, our results are ultimately constrained by the scale of brain activity accessible by the 1.8-mm resolution fMRI protocol and the modality’s inability to capture temporal dynamics of language processing. Second, the meaning of some of the six-word-long sentences might be underdetermined, and may therefore lead to different representations across individuals. For example, two people hearing “As if that would impress anyone” might imagine highly distinct situations where such a sentence could be uttered (Hosseini et al. 2024). Third, although we chose sentences that span diverse topics and styles, no set of 200 sentences covers the full space of linguistic variation, so a larger and more diverse set may reveal additional components. Fourth, given that our participants varied in their linguistic and cultural backgrounds (e.g., monolingual English speakers and multilingual individuals who acquired English during childhood), a more homogeneous participant pool might have uncovered additional shared components.

How do the current findings relate to prior studies that have found ‘semantic tuning’—a preference for particular kinds of meanings—during narrative comprehension (Huth et al. 2016; Deniz et al. 2019)? These previous efforts characterize voxel-level tuning during language comprehension using decomposition-based approaches, and show that extensive portions of association cortex contain voxels tuned to broad semantic categories related to e.g., social, tactile, or numeric content. Superficially, these results appear inconsistent with our findings: the cortical regions driven by our experiment are more spatially restricted, and we do not find any shared components that relate to particular kinds of meanings, except for the general preference for more abstract meanings. We speculate that the differences are due to differences in the study design and analytic approaches. First, Huth et al.’s (2016) results are strongly tied to the choice of model: voxel responses were modeled using a 985-parameter semantic feature space based on word co-occurrence statistics, and principal components were derived from the resulting regression weights. This model-first approach constrains the findings to dimensions captured by the chosen model space (word co-occurrence statistics) and may not capture other dimensions (e.g., properties related to linguistic form). Huth et al.’s test for shared dimensions was conducted within the space of their semantic model, evaluating whether dimensions derived from *n*-1 participants could predict the model regression weights of the held-out participant. As a result, the shared dimensions were defined relative to the semantic features defined by the model. In contrast, our approach is data-driven and model-free: we apply dimensionality reduction directly to voxel responses and test for generalization of voxel-derived components without assuming a predefined model space. Second, Huth et al. (2016) and Deniz et al. (2019) used continuous narratives, whereas we used short, decontextualized sentences. Processing the long contexts inherent in narratives likely engages brain areas outside of the fronto-temporal language network, such as the ‘Default Mode Network’ (Regev et al. 2013; Yeshurun et al. 2017; Nguyen et al. 2019; Blank & Fedorenko 2020), which may explain the greater spatial extent of the effects observed in those studies. Note that in narratives, it may be difficult to determine whether semantic tuning reflects word-, phrase-, or discourse-level processes (Rust & Movshon 2005; Hamilton & Huth 2018). On the whole, we believe that the results across studies are not necessarily inconsistent, but rather reflect different research questions and analytic approaches, as well as reliance on paradigms that probe distinct aspects of language comprehension and semantic processing. To better understand how the findings relate to one another, future work could bridge these paradigms and directly compare model-based and model-free approaches.

### 2. Organizing dimensions of the fronto-temporal language network

The language network, a set of frontal and temporal regions in the left hemisphere, has been widely studied and characterized as a modality-independent network selectively and causally supporting language processing (Fedorenko et al. 2024). These regions are typically identified using a short localizer experiment contrasting brain responses to well-formed linguistic inputs versus a perceptually matched control condition (Fedorenko et al. 2010; Scott et al. 2017; Tuckute, Lee et al. 2024c), but can also be recovered from task-free fMRI data from voxel-wise patterns of functional correlations (Braga et al. 2020; Du et al. 2024; Shain & Fedorenko 2025). Prior studies have shown that the five regions that constitute this “language network” exhibit similar functional response profiles (e.g., Rodd et al. 2010; Fedorenko et al. 2020; Hu et al. 2022; Shain, Blank et al. 2020; Wehbe et al. 2021), leaving the internal organization of this network elusive. To address this gap, we searched for the organizing dimensions of the language network by examining voxel-level responses to diverse sentences, leveraging the high resolution and high sensitivity of 7T fMRI.

In line with past evidence of similar responses across the five language fROIs, we found that the two Sentence PCs—processing difficulty and meaning abstractness—were present in every fROI, with every region showing stronger response to sentences that are hard to process and express abstract meanings (see also Willems et al. 2016; Henderson et al. 2016; Shain, Blank et al. 2020; Wehbe et al. 2021; Tuckute et al. 2024d; Botch & Finn 2024). Very few voxels within these language fROIs showed a preference for easy-to-process/concrete sentences (only 8% of fROI voxels could be categorized as easy-to-process or concrete-preferring; <u>SI 11</u> and Figure 5B). Notably, although the language fROIs showed an overall similar tuning profile (i.e., the mean angle is roughly similar across fROIs; <u>SI 12</u> and Figure 5B), the internal *composition* of voxel tuning profiles varied among fROIs: for instance, the anterior temporal fROI contains a large proportion of voxels preferring abstract sentences (e.g., Pobric et al. 2007; Simmons & Martin, 2009) whereas the IFG fROI does not (<u>SI 11</u> and Figure 5B). In general, the two temporal fROIs contained a larger proportion of abstract-preferring voxels compared to the frontal ones (<u>SI 11</u>). These results demonstrate that there is meaningful voxel-level variation across and within the language fROIs—variation that may be important for supporting distinct aspects of linguistic computation. More broadly, our findings highlight the importance of examining cortical tuning at the level of individual voxels.

When we expanded our analysis beyond the individually defined language fROIs containing the most language-selective voxels to include nearby voxels within the anatomical language parcels, we observed an overall similar tuning profile to those in the fROIs, but with a more diverse composition of tuning properties (i.e., a broader span of the rainbow color wheel; compare the two panels in Figure 5). Importantly, the *differences* among language parcels were largely driven by voxels that are not language-selective (see <u>Results; How are the components spatially organized?</u>). Together, these findings suggest that the fROIs, defined through the localizer contrast, carve out a specific part of the tuning landscape, whereas nearby less/or not language-selective voxels respond reliably to sentences but vary more in their sentence tuning. We speculate that these non-language selective voxels with more diverse tuning profiles might facilitate interactions between the language regions and other networks in the brain.

### 3. Voxels with preference for concrete sentences in the ventral temporal cortex

The two Sentence PC model showed robust predictivity in the lateral frontal and temporal brain areas traditionally associated with language processing (Binder et al. 1997; Lipkin et al. 2022), but also predicted responses in voxels located in the left ventral temporal cortex (VTC), primarily associated with high-level vision (Figure 4 and <u>SI 3</u>, <u>SI 10</u>). During the experiment, the sentences were presented auditorily and participants were not instructed to perform any visual tasks; they were asked to listen attentively and think about the sentence’s meaning, and later complete a memory task. Yet, these VTC voxels were significantly predicted by the two PCs and exhibited a preference for concrete sentences—a tuning profile that differed from the fronto-temporal language network.

What could explain reliable sentence responses in the VTC? One possibility is that these responses reflect visual imagery. The VTC is known for its role in high-level visual processing, including category-selective areas involved in recognizing faces, bodies, and places (Kanwisher et al. 1997; Epstein & Kanwisher 1998; Peelen & Downing 2005; Grill-Spector & Weiner 2014). Critically, these and nearby areas have been shown to be active not only during perception, but also during imagery of (O’Craven & Kanwisher 2000; Ganis et al. 2004; Lee et al. 2012) and memory for these categories (Bainbridge et al. 2021; Steel et al. 2023). We examined voxel-level angle tuning in these category-selective regions (Rosenke et al. 2021) and found that the well-predicted areas were in or proximal to the fusiform-face area 2 (FFA-2), fusiform-body area (FBA), and parahippocampal place area (PPA) (<u>SI 10</u>), which could have been activated through visual imagery of concrete sentences.

Another possibility is that the responses originate not in the category-selective visual areas, but in a language area in the VTC. In particular, a “basal temporal language area (BTLA)” has been proposed as a multimodal area that processes both language and visual information (Lüders et al. 1991; Purcell et al. 2014; Li et al. 2024). Recently, Li et al. (2024) characterized two language-responsive areas within the left VTC (see Wolna et al. (2025) for concordant evidence using a similar functional localization approach). One area, located anteriorly, close to the temporal pole, responded significantly more strongly to sentences than all visual categories, whereas the second area, medially located, showed comparable response to auditory language and visual conditions. Our well-predicted voxels overlapped with the medial area identified by Li et al. (2024), with these voxels exhibiting a preference for concrete sentences (<u>SI 10</u>). The medial language parcel partly overlapped with some of the visual parcels (FFA-2 and FBA), though the spatial extent of our well-predicted voxels went beyond the estimated boundaries of this medial language area (<u>SI 10</u>). More generally, if (some of) these voxels constitute a multimodal language area in the VTC, our findings suggest that their computations differ from those implemented in the core frontal and temporal language areas.

Further work is needed to understand the processes taking place in the well-predicted VTC regions. A limitation of the current study is the lack of localizer experiments targeting imagery or other functions typically associated with the ventral stream (e.g., Kanwisher et al. 1997; Downing et al. 2001; Cohen et al. 2002). Nevertheless, we emphasize that the two Sentence PCs can explain activity in regions beyond typical fronto-temporal language network, and their predictions appear meaningfully situated within the broader topography of the brain—with tuning for concrete sentences in areas associated with high-level vision.

### 4. What are the two Sentence PCs representing and computing?

Our findings reveal two shared components in language processing. What is the most accurate characterization of these components, and what might be their functional role in language processing? We characterized the two PCs using a set of 16 linguistic/semantic properties. Sentence PC 1 was strongly correlated with measures of linguistic processing difficulty. As a direct behavioral measure, we collected reading times using the Maze task (Forster et al. 2009; Freedman & Forster, 1985), where participants incrementally choose, at each position in a sentence, between the correct next word and a distractor word. These word-by-word response times are widely used in behavioral measurements of language comprehension and give relatively direct insight into processing difficulty. We extended the approach from Boyce et al. (2020; 2023), who automated distractor selection by identifying high-surprisal words using language models, by leveraging a more recent language model (GPT2; Radford et al. 2018) to generate well-matched context-sensitive distractors (<u>SI 13</u>). Sentence PC 1 was highly correlated with the average sentence reading time: sentences with high loadings on this PC take longer to read, establishing a link between neural response tuning and behavioral processing difficulty. We also tested a broader set of linguistic properties that have been shown to affect language comprehension. Psycholinguistic studies have shown that sentences using low-frequency words and constructions and higher-surprisal sentences are harder to process (e.g., Demberg & Keller 2008; Smith & Levy 2013; Singh et al. 2016; Brothers & Kuperberg 2021; Hahn et al. 2022). Motivated by these findings, we tested three different surprisal measures, pointwise mutual information (PMI, capturing local co-occurrence structure), and behavioral ratings of whether sentences are perceived as grammatical, plausible, and frequent. Sentences with higher PC 1 loadings were more surprising, had lower PMI, and were rated as less well-formed, less plausible, and less frequent. These results reinforce the interpretation of Sentence PC 1 as reflecting a processing difficulty axis. This PC likely reflects multiple aspects of linguistic processing difficulty: evidence from psycholinguistics and neuroscience has demonstrated processing costs associated with predicting upcoming words (Hale 2001; Levy 2008; Lopopolo et al. 2017; Shain, Blank et al. 2020; Heilbron et al. 2022) as well as costs from storing and integrating words into the evolving representation of the sentence in memory (Gibson 2000; Ben-Shachar et al. 2003; Constable et al. 2004; Blank et al. 2016; Shain, Kean et al. 2024a). Disentangling these contributions is challenging (but see Shain et al. (2022)), particularly given the relatively short sentences (six words) used in our study. Hence, Sentence PC 1 may reflect fundamental linguistic computations such as prediction and integration during sentence comprehension, though it may also reflect additional, yet unidentified, computations.

Sentence PC 2 showed a distinct profile. This PC was strongly correlated with behavioral ratings of concreteness—how much a sentence’s meaning is tied to perceptual experience, including visual experience. Sentences with high loadings on this PC are hard-to-visualize and abstract. Thus, our candidate explanation for this is meaning abstractness. However, we note that some form-based features covary with abstractness—for instance, concrete words tend to have a larger proportion of content words (nouns, verbs, adjectives and adverbs) (*r* = 0.12), so sentences rated as more abstract contain a higher proportion of function words, such as prepositions, determiners, and pronouns. Accordingly, whereas concrete sentences might convey much of their meaning through the semantics of individual content words, abstract sentences may rely more on meaning that emerges from semantic compositional computations.

Decades of theoretical and behavioral research have emphasized the importance of the concrete-to-abstract axis in organizing mental representations (e.g., Paivio, Allan 1971; Pecher et al. 2011; Borghi et al. 2017). Neuroscience studies have examined brain responses to concrete and abstract language, largely focused on single words (West & Holcomb 2000; Binder et al. 2005; Moseley & Pulvermüller 2014; Roxbury et al. 2014; Fernandino et al. 2015; Wang & Bi 2021; cf. Botch & Finn 2024). One key finding is that neural representations of concrete words appear to be more similar across individuals, and also more similar within the same individual across distinct exposures (Wang & Bi 2021; Botch & Finn 2024). In the current work, we establish that the abstractness of a sentence modulates brain responses during sentence comprehension, and show that this dimension is shared across individuals.

What does this study, and the emergence of the two shared components, tell us about how meaning is extracted from language? We believe that the components reflect a robust, low-dimensional subspace in the frontal and temporal language areas that likely serve as the representational scaffold for meaning extraction. For example, signaling that a sentence is abstract may constrain and thus facilitate access to the relevant parts of broader knowledge representations distributed across the brain (e.g., Huth et al. 2016), which might be unique to each individual. To test such hypotheses and to probe the functional roles of the two shared components, artificial neural network models for language will be invaluable (Jain et al. 2023; Tuckute et al. 2024a,b) due to the limited tools available to probe human neural representations of language. These models may provide insights into how these two components work together, to shed light on the mechanisms underlying language comprehension in the human brain.

## Data and code availability

Data and code will be made publicly available upon publication.

## Acknowledgements

We thank Emily Allen for help with fMRI scanning. We are grateful for feedback on the project from Jacob S. Prince and Nicholas M. Blauch. We thank Selena She, Maya Taliaferro, and Thomas Clark for help with linguistic properties. We thank Jin Li for providing the language VTC parcel. Greta Tuckute was supported by a Friends of McGovern Graduate Fellowship and the K. Lisa Yang ICoN Center Graduate Fellowship. Ev Fedorenko was supported by research funds from the McGovern Institute for Brain Research, the Department of Brain and Cognitive Sciences, MIT’s Quest for Intelligence, and a grant from the Simons Foundation to the Simons Center for the Social Brain at MIT. Kendrick Kay acknowledges funding from National Institutes of Health grant (NIH) R01EY034118.

## Methods

### 1. fMRI experiments

#### Participants

Eight participants (six males and two females; age range 18–42 years; one is a co-author) took part in this study between December 2023 and May 2024. All participants had normal or corrected-to-normal visual acuity and no history of neurological, developmental, or language impairments. All participants were right-handed, as determined by self-report and the Edinburgh handedness inventory (Oldfield 1971). All participants had native-like proficiency in both spoken and written English for both comprehension and production, as determined by self-report. Two participants were native English monolingual speakers (P7, P8). Three participants were native English speakers; of these, two had proficiency in one additional language (P2, P3), and one had proficiency in two additional languages (P6). The remaining three participants were not native English speakers, but were exposed to English before age six and had proficiency in two additional languages. Informed written consent was obtained from all participants, and the experimental protocol (1508M77463) was approved by the University of Minnesota Institutional Review Board. Participants were compensated at a rate of $30 per hour, plus an additional performance bonus. All participants had high data quality, and no participants were excluded from this study. Each participant took part in one fMRI scan session that lasted around three hours. Participants had the option to take a break halfway through the session.

#### Sentence-listening experiment

##### Stimulus selection and stimulus preprocessing

We selected a set of 200 sentences to be presented three times each within a single, extended scanning session. We first took the set of 1,000 “baseline” sentences from Tuckute et al. (2024d). In brief, these 1,000 sentences were sampled from naturalistic corpora in order to achieve a stylistically and semantically diverse set. Specifically, sentences were chosen to span a range of sentence styles as well as a large range of topics based on semantic clustering as quantified through GloVe vectors (Pennington et al. 2014). In Tuckute et al. (2024d), the sentences were visually presented to the participants. To select sentences for auditory presentation in the current study, sentences were not sub-selected based on sentence content or other linguistic criteria; instead, sentences were selected based on a combination of synthesized audio quality and random sampling. The selection process proceeded in five main steps, as detailed below.

*First*, we used the Google Cloud Text-to-Speech service with the “Studio_O” voice identity to generate auditory versions of the 1,000 sentences from Tuckute et al. (2024d). With an initial starting speech rate of 1.0, we iteratively re-generated an audio clip with an adjusted speech rate such that the generated audio clip is 2 seconds long. The speech rate was iteratively adjusted, constrained to values between 0.5 and 1.5. Sentences that did not meet this duration criterion were excluded, leaving 854 sentences. *Second*, we excluded sentences containing numbers above 20 (2 sentences), sentences with abbreviations (13 sentences), and sentences with characters other than A-Z, 0-9, periods, commas, exclamation marks, question marks, apostrophes, and quotation marks (34 sentences). This step left 805 sentences. *Third*, we further constrained the sentence set by limiting the speech rate to values between 0.75 and 1.25, leaving 478 sentences. *Fourth*, from these 478 sentences, 300 were randomly sampled and manually reviewed by an experimenter for quality. Sentences with artifacts, abnormal speech rates, or poor intonation were excluded. Based on this screening, 200 final sentences were selected. *Finally*, the audio samples were preprocessed: We added a 25-ms linear in and out ramp to avoid clicking artifacts, resampled the audio samples from 24kHz to 44.1kHz, and normalized the volume of each audio clip to an RMS of 0.1.

##### fMRI design

Participants listened to each sentence in an event-related fMRI design. Each sentence item was presented three times for a total of 200 sentence items × 3 presentations: 600 trials. Sentences were presented auditorily for 2 s followed by a 4-s inter-stimulus interval. Throughout the experiment, the screen displayed a black fixation cross on a gray background. Participants were instructed to listen attentively and think about the meaning of the sentences while maintaining fixation on the fixation cross. To encourage engagement with the stimuli, participants were informed that they would be asked to perform a short memory task after the session (outside of the scanner) and that the memory task would be conducted in a different modality (text).

Each run contained 50 unique sentence trials and four 10-s fixation blocks (in the beginning, after the 17th and 33rd sentence trials, and at the end of each run). Each run lasted 340 s (5:40 minutes) and participants completed 12 runs in total.

Sentence items were pseudorandomly distributed across runs. For each participant, the first repetition of each item was presented in the first four runs (in random order), the second repetition of each item was presented in the next four runs (in a different random order; runs 5–8), and the third and final repetition was presented in the last four runs (in yet another different random order; runs 9–12). To minimize contextual effects, item order was randomized for each repetition and independently for each participant.

##### Behavioral assessment for the sentence-listening experiment

Participants completed a memory task at the end of the scanning session (outside of the scanner) as a behavioral assessment of their engagement during the sentence-listening experiment. Participants were visually presented with sentences, one at a time, and asked to decide whether they had heard each sentence during the sentence-listening experiment. The memory task consisted of 30 unique sentences: 15 target sentences from the set presented in the scanning session and 15 foil sentences. For target sentences, we randomly selected one sentence from each of the 12 fMRI runs and randomly selected three additional sentences for a total of 15 unique target sentences. For foil sentences, we randomly sampled without replacement from a set of 60 six-word-long sentences taken from nine different text corpora (consisting of published written text, web media text, and transcribed spoken text; see <u>SI 1</u> in Tuckute et al. (2024d)). Importantly, the corpora from which the foils were drawn were also used to construct the stimulus set for the sentence-listening experiment, ensuring stylistic similarity.

The average accuracy on the memory task (sum of correct responses divided by the total number of responses; chance level is 50%) was 91.7% (SD across participants = 8.2%; min = 73.3%; max = 100%).

#### Language network localizer experiment

The task used to localize the language network was a reading task contrasting sentences (e.g., THE SPEECH THAT THE POLITICIAN PREPARED WAS TOO LONG FOR THE MEETING) and lists of unconnected, pronounceable nonwords (e.g., LAS TUPING CUSARISTS FICK PRELL PRONT CRE POME VILLPA OLP WORNETIST CHO) in a standard blocked design with a counterbalanced condition order across runs (introduced in Fedorenko et al. (2010)). The *sentences > nonwords* contrast targets high-level aspects of language, including lexical and phrasal semantics, morphosyntax, and sentence-level pragmatic processing, to the exclusion of perceptual (speech- or reading-related) processes. The areas identified by this contrast are strongly selective for language relative to diverse non-linguistic tasks (see Fedorenko & Blank (2020); and Fedorenko et al. 2024 for reviews). This paradigm has been extensively validated and shown to be robust to variation in the materials, modality of presentation, language, and task (e.g., Malik-Moraleda et al. 2022; Scott et al. 2017; Tuckute, Lee, et al. 2024c). Further, a network that corresponds closely to the localizer contrast emerges robustly from whole-brain resting-state (task-free) data (Braga et al. 2020).

Each stimulus consisted of 12 words/nonwords. Stimuli were presented in the center of the screen, one word/nonword at a time, at the rate of 450 ms per word/nonword. Each stimulus was preceded by a 100-ms blank screen and followed by a 400-ms screen showing a picture of a finger pressing a button and another 100-ms blank screen, for a total trial duration of 6 s. Stimulus blocks lasted 18 s (3 trials per block), and fixation blocks lasted 14 s. Each run (consisting of 5 fixation blocks and 16 stimulus blocks) lasted 358 s (5:58 min). Participants completed 2 runs. Participants were instructed to read attentively (silently) and press a button on the button box whenever they saw the finger picture. The button-pressing task was included to help participants remain alert. Experiment materials and scripts are available from the Fedorenko Lab website (https://evlab.mit.edu/funcloc).

#### MRI acquisition

##### MRI data acquisition

MRI data were collected at the Center for Magnetic Resonance Research at the University of Minnesota. Anatomical (T1- and T2-weighted) data were collected with a 3T Siemens Prisma scanner and a standard Siemens 32-channel RF head coil. Functional data (T2*-weighted) were collected using a 7T Siemens Magnetom actively-shielded scanner and a single-channel-transmit, 32-channel-receive RF head coil (Nova Medical, Wilmington, MA). To mitigate head motion, we used standard MR-compatible foam padding on the back of the head, along with additional neck and ear padding.

We collected anatomical data at 3T. The motivation for collecting anatomical data at 3T was to ensure acquisition of T1-weighted volumes with good gray/white-matter contrast and good spatial homogeneity, which is difficult to achieve at ultra-high fields. To increase contrast-to-noise ratio, we acquired several repetitions of T1- and T2-weighted volumes. For each participant, we collected between 1–9 scans (median: 3) of a whole-brain T1-weighted MPRAGE sequence (0.8-mm isotropic resolution, repetition time (TR) 2400 ms, echo time (TE) 2.22 ms, TI 1000 ms, flip angle 8°, bandwidth 220 Hz/pixel, no partial Fourier, in-plane acceleration factor (iPAT) 2, time of acquisition (TA) 6.6 min/scan) and 1–3 (median: 2) scans of a whole-brain T2-weighted SPACE sequence (0.8-mm isotropic resolution, TR 3200 ms, TE 563 ms, bandwidth 744 Hz/pixel, no partial Fourier, iPAT 2, TA 6.0 min/scan).

We collected functional data at 7T. Functional data were collected using gradient-echo EPI at 1.8-mm isotropic resolution with whole-brain (including cerebellum) coverage (84 axial slices, slice thickness 1.8 mm, slice gap 0 mm, field-of-view 216 mm (FE) x 216 mm (PE), phase-encode direction anterior-to-posterior, matrix size 120 x 120, TR 1600 ms, TE 22.0 ms, flip angle 62°, echo spacing 0.66 ms, bandwidth 1736 Hz/pixel, partial Fourier 7/8, iPAT 2, multiband slice acceleration factor 3). In addition to the EPI scans, we also collected dual-echo fieldmaps for post-hoc correction of EPI spatial distortion (same overall slice slab as the EPI data, 2.2 mm x 2.2 mm x 3.6 mm resolution, 42 slices, TR 510 ms, TE1 8.16 ms, TE2 9.18 ms, flip angle 40°, bandwidth 350 Hz/pixel, partial Fourier 6/8, TA 1.3 min/scan). Fieldmaps were periodically acquired over the course of each scan session to track changes in the magnetic field.

For participant 6, the scanner stopped prematurely during run 11 of the 12 sentence-listening experiment runs, missing the final 10 volumes (less than 5% of the run). To alleviate this problem, the missing volumes were simply replaced with the last acquired volume.

##### Stimulus presentation

Visual stimuli were presented using a Cambridge Research Systems BOLDscreen 32 LCD monitor positioned at the head of the 7T scanner bed, placed flush against the scanner bore. The monitor operated at a resolution of 1920 pixels × 1080 pixels at 120 Hz. The size of the full monitor image was 69.84 cm (width) × 39.29 cm (height). Participants viewed the monitor via a mirror mounted on the RF coil. The viewing distance was ∼5.5 cm from the participants’ eyes to the mirror + 178.5 cm from the mirror to the monitor image = ∼184 cm total. Auditory stimuli were presented via MRI-compatible Sensimetrics S15 foam tip earbuds with filters to flatten the frequency response. Before the main sentence-listening experiment, each participant performed volume adjustment during the acquisition of a mock functional run. During this run, speech sounds—similar but not identical to those in the critical experiment—were played, and participants were asked to press a button to indicate whether the volume should be louder or quieter, or whether the volume is just right, with the goal of being comfortable and audible over the scanner noise. The adjusted volume was then used for the entire experimental session. Behavioral responses were recorded using a button box (Current Designs).

##### Eyetracking

Eyetracking was performed using an EyeLink 1000 system (SR Research, Mississauga, Ontario, Canada) combined with a custom infrared illuminator mounted on the RF coil. Based on the eyetracking, we continuously monitored participants’ alertness throughout the experimental session, but actual eyetracking data were not saved. In a few instances, drowsy participants were encouraged to take short breaks before proceeding with the remainder of the experiment.

#### MRI preprocessing and first-level analysis

##### MRI preprocessing

The data were preprocessed using a custom pipeline (based on methods adapted from Kay et al. (2019) and Allen et al. (2022)). Preprocessing results were carefully visually inspected to ensure quality control.

##### T1- and T2-weighted volumes

T1- and T2-weighted volumes were corrected for gradient nonlinearities using a custom Python script (https://github.com/Washington-University/gradunwarp) and the proprietary Siemens gradient coefficient file retrieved from the scanner. The T1 (or T2) volumes acquired for a given participant were then co-registered. This was accomplished by co-registering all T1 (or T2) volumes (rigid-body transformation; correlation cost metric). In the estimation of registration parameters, a manually defined 3D ellipse was used to focus the cost metric on brain tissue. Multiple acquired volumes were cubic interpolated to implement the co-registration, and then averaged to increase contrast-to-noise ratio. This resulted in a single averaged T1-weighted volume and a single averaged T2-weighted volume.

The averaged T1-weighted volume was processed using FreeSurfer version 7.4.1 with the -hires option (to take advantage of the 0.8-mm resolution). To improve cortical surface reconstruction accuracy, we also provided the averaged T2-weighted volume to FreeSurfer, and specified the -T2pial option. As part of FreeSurfer’s processing, the T2-weighted volume was aligned and resampled to match the T1-weighted volume. Using mris_expand, we generated cortical surfaces positioned at 10%, 26%, 42%, 58%, 74%, and 90% of the distance between the pial surface and the boundary between gray and white matter.

##### fMRI preprocessing

In the first stage of fMRI preprocessing, the fMRI data were preprocessed by performing temporal resampling (cubic interpolation) to correct for slice time differences and to also upsample the data (in the same step) to 1.0 s. Fieldmaps were linearly interpolated over time, producing an estimate of the field for each fMRI volume acquired, and then temporally resampled fMRI volumes were undistorted based on the field estimates. The undistorted volumes were then used to estimate rigid-body motion parameters using the SPM5 utility spm_realign (a manually defined 3D ellipse was used to focus the cost metric on brain regions unaffected by gross susceptibility effects). Finally, a spatial resampling (cubic interpolation) on the temporally resampled volumes was performed to correct for the combined effects of head motion and EPI distortion. In the second stage, we aligned the mean preprocessed fMRI volume to the T2-weighted volume (prepared by FreeSurfer) using a small amount of nonlinear warp as determined using ANTs 2.1.0 (Avants et al. 2009). To maximize registration quality, an independent registration was performed for the mean volume associated with each fMRI run. The parameters of the ANTs registration were tuned to maximize registration accuracy (while avoiding overfitting). In the third stage, we coupled the nonlinear warp estimated in the second stage with the first-stage transformations to produce final preprocessed fMRI volumes in register with the anatomy. For this step, we used a 1.8-mm voxel grid parallel to the FreeSurfer anatomical volume. The final result of fMRI preprocessing was volumetric fMRI time-series data in subject-native space (registered to the anatomy).

##### fMRI first-level analysis

Data were analyzed using GLMsingle (Prince et al. 2022) using default parameters. The output of GLMsingle consisted of single-trial response amplitude (i.e. betas) in units of percent signal change in subject-native volumetric space. The betas were mapped using cubic interpolation to the four middle cortical surfaces positioned at 26%, 42%, 58%, and 74% of the distance between the pial surface and the boundary between gray and white matter. Next, these four sets of betas were averaged across surface depth, and then the averaged betas were transformed to the fsaverage cortical group surface (Fischl et al. 1999) using nearest neighbor interpolation (163,842 vertices per hemisphere). Note that all analyses were conducted in the fsaverage surface space, but for simplicity we refer to vertices as “voxels” throughout this paper.

### 2. Definition of regions of interest (ROIs)

#### Anatomical ROIs

##### Language parcels

The language parcels were derived using watershed parcellation based on a probabilistic activation overlap map, as described by Fedorenko et al. (2010) for the contrast of *sentences > nonwords*. The dataset used for this procedure consisted of 220 independent participants (the parcels are available at https://www.evlab.mit.edu/resources-all/download-parcels). Parcels covered extensive portions of the lateral frontal, temporal, and parietal cortices of the left hemisphere. Specifically, there were five parcels: three on the lateral surface of the frontal cortex (in the inferior frontal gyrus, *IFG*, and its orbital part, *IFGorb*, as well as in the middle frontal gyrus, *MFG*), and two on the lateral surface of the temporal and parietal cortex (in the anterior temporal cortex, *AntTemp*, and posterior temporal cortex, *PostTemp*). The parcels were defined in the Montreal Neurological Institute (MNI) template space (XI549Space) and were projected to the fsaverage space using the NeuroMaps software (Markello et al. 2022) using *k*-nearest neighbors interpolation. To eliminate interpolation artifacts, we manually performed small corrections to the parcels such that they were contiguous parcels on the fsaverage surface. To ensure that the parcels cover language-selective cortex in our participants, we visually inspected the parcels overlaid on the results of the language localizer experiment conducted in this study. Visualizing the *t*-statistic averaged across participants (*sentences > nonwords*, <u>Methods; Language</u> <u>network localizer experiment</u>), we noted that the three frontal parcels did not sufficiently encompass the localizer activity. To address this, we manually expanded the frontal parcels. Finally, following prior work (e.g., Blank et al. (2014)), to define parcels in the right hemisphere, the parcels in the left hemisphere were mirrored onto the right hemisphere using FreeSurfer’s xhemi surface.

##### Visual parcels

The visual parcels were derived from a probabilistic activation map from experiments targeting early visual and category-selective regions in human ventral and lateral occipito-temporal cortex as described in Rosenke et al. (2021). The atlas (available at <u>download.brainvoyager.com/data/visfAtlas.zip</u>) consists of 17 parcels in each hemisphere, with 11 of these parcels being category-selective ones: mFus_faces, pFus_faces, IOG_faces, OTS_bodies, ITG_bodies, MTG_bodies, LOS_bodies, pOTS_characters, IOS_characters, CoS_places, and hMT_motion. The remaining six parcels are retinotopic, which we did not include in our analyses. We further excluded MTG_bodies because it overlapped with the anatomical PostTemp language parcel (453 voxels), leaving 10 parcels of interest.

##### Ventral temporal language parcel

The language parcel in the ventral temporal cortex (VTC) was derived from a probabilistic activation overlap map, as described by Li et al. (2024) for the contrast of *sentences > texturized sentences* (auditorily presented). The atlas (available at https://www.zeynepsaygin.com/ZlabResources.html) consists of two parcels in the left hemisphere, an anterior VTC language parcel and a medial VTC language parcel. We included the medial parcel as it showed overlap with the language-evoked responses in the current study.

#### Functional ROIs

##### Language network fROIs

For each participant, functional regions of interest (fROIs) were defined by combining two sources of information: i) the participant’s activation map for the localizer contrast of interest (*t*-map), and ii) group-level constraints as given by the anatomical parcels described above. These anatomical parcels delineate the expected gross locations of activations for the contrast and were sufficiently large to encompass the extent of variability in the locations of individual activations.

The language fROIs were defined using the *sentences > nonwords* contrast from the language localizer (Fedorenko et al. 2010) (<u>Methods; Language network localizer experiment)</u>. This contrast targets higher-level aspects of language, to the exclusion of perceptual (speech/reading) and motor-articulatory processes (for discussion, see Fedorenko & Thompson-Schill, 2014). To define the language fROIs, each participant’s *sentences > nonwords t*-map was intersected with a set of five binary parcels in the left hemisphere (described in <u>Methods; Language</u> <u>parcels</u>). Within each of these five parcels, the 10% of voxels with the highest *t*-values for the *sentences > nonwords* contrast were selected.

### 3. Noise ceiling estimation

Noise ceilings were estimated to obtain a measure of data reliability. The noise ceiling for a given voxel is defined as the maximum percentage of variance in the voxel’s responses that in theory can be explained, given the presence of noise (defined as response variability across repetitions of an item). The procedure for noise ceiling estimation was similar to that of Allen et al. (2022).

First, we computed the variance of the betas across the three presentations of each sentence (using the unbiased estimator which normalizes by *n*-1 where *n* is the number of trial repetitions), averaged this variance across sentences and then computed the square root of the result. This step provides an estimate of the noise standard deviation:

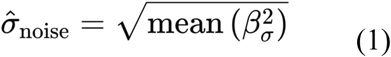

where 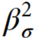 indicates the variance across the betas obtained for a given sentence. Second, we computed the variance of the trial-averaged sentence data 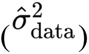, and the signal standard deviation was estimated as:

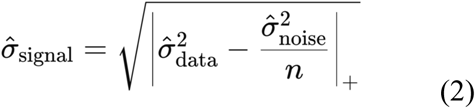

where ||+ indicates positive half-wave rectification. Note that the noise variance is divided by the number of trials (*n* = 3) because data variance is computed on the trial-averaged data. Third, the noise ceiling signal-to-noise (NCSNR) was computed as the ratio between the signal and noise standard deviations:

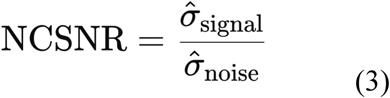

The NCSNR metric was used to guide inclusion of voxels in the derivation of Sentence PCs (<u>Methods; Deriving Sentence PCs</u>). Finally, the noise ceiling (NC) was then computed as the amount of variance contributed by the signal expressed as a percentage of the total amount of variance in the data:

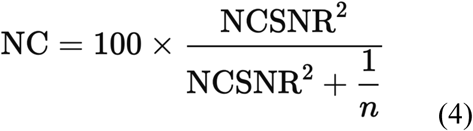

Noise ceilings were used in Figure 2 to evaluate the performance of the Sentence PC models with respect to the theoretical upper bound of explainable variance in the voxel responses.

### 4. Deriving Sentence PCs

Sentence PCs were derived through matrix decomposition of a data matrix (**Z**) consisting of beta weights in percent signal change. Our procedure consisted of the following steps.

#### Data preparation

We started with a data matrix **Z** with dimensionality sentences × voxels × trials. We averaged responses across trials, yielding trial-averaged data **Z_avg_** with dimensionality sentences × voxels. We subtracted each voxel’s mean response from the trial-averaged data, yielding mean-subtracted trial-averaged data **Z_ms_** with dimensionality sentences × voxels.

Voxels in **Z** were extracted from five anatomical parcels that broadly encompass the language network (<u>Methods; Language parcels</u>). Voxels were pooled across parcels and participants. To analyze reliable voxels, voxels were selected based on a voxel inclusion criterion, and to reduce computational load, these voxels were further randomly subsampled. The primary analyses in this paper used the following criteria and subsampling parameters: a noise ceiling signal-to-noise ratio (NCSNR) threshold of >0.4 within language parcels and 75% of these voxels randomly subsampled due to computational load. These parameters yield 14,613 voxels across eight participants. Sentence PCs were robust to different voxel inclusion parameters (<u>SI 1</u>).

#### Calculation of voxel covariance

We proceeded to obtain a voxel covariance matrix, which serves to estimate the main axes of variance in the data. We obtained the voxel covariance matrix using two different approaches:

i. Our primary approach leverages a technique called Generative Modeling of Signal and Noise (GSN; Kay et al. 2024), which explicitly characterizes neural responses in terms of signal (responses that are reliably driven by each experimental condition) and noise (trial-to-trial variability in responses for the same condition). Specifically, GSN involves modeling each data point in **Z** as the sum of a sample from a signal distribution and a sample from a noise distribution. By applying GSN to **Z**, we obtained two voxel covariance matrices, **C_signal_** and **C_noise_**, each with dimensionality voxels × voxels. The advantage of the GSN approach is that the influence of noise on voxel covariance is isolated and removed.
ii. As a supplementary analysis, we also implemented a naive approach in which we simply calculate the covariance of **Z_ms_**, yielding a voxel covariance matrix **C_naive_** with dimensionality voxels × voxels. Note that in contrast to the GSN approach, **C_naive_** is affected by both signal and noise.

#### Calculation of eigenvectors via SVD

We performed singular value decomposition (SVD) on **C_signal_** (approach i) or **C_naive_** (approach ii). In both cases, we obtained a decomposition of the voxel covariance matrix as **USV^T^**. **V** is an orthonormal matrix with dimensionality voxels × voxels, and consists of unit-length eigenvectors in the columns. **S** is a diagonal matrix with dimensionality voxels × voxels, and has eigenvalues in descending order along the diagonal. **U** is identical to **V**. As a pre-processing step, given that the sign of each eigenvector is arbitrary, we checked the mean of each eigenvector and if the mean was less than zero, we flipped the sign for that eigenvector. Note that the sum of the first *n* eigenvalues expressed as a percentage of the sum of all eigenvalues is the amount of variance accounted for by the first *n* eigenvectors.

Intuitively, the eigenvectors define a new basis in which to express the data. The eigenvectors are ordered, and each subsequent eigenvector denotes a direction in voxel space that accounts for the maximal amount of variance in the data.

#### Derivation of Sentence PCs

To determine how much each sentence falls along the axes of variation as defined by the eigenvectors, we took the mean-subtracted data and projected them onto the eigenvectors: **score** = **Z_ms_V**. **score** is a matrix of scores with dimensionality sentences × eigenvectors, where the *i*th column indicates the coordinates of each sentence response along the *i*th eigenvector. We refer to the columns of **score** as ‘Sentence PCs’ throughout the paper. Intuitively, each Sentence PC indicates how different sentences drive voxel responses (across language regions and participants). We interpret the derived Sentence PCs as a set of response components that can be combined to characterize sentence responses from any given voxel (see next section). In a supplementary analysis, we explored using non-negative matrix factorization as an alternative approach to deriving response components (<u>SI 1</u>).

Note that for simplicity, we interpret Sentence PCs as potentially having one-to-one correspondence with sentence properties (<u>Methods; Sentence properties</u>). However, there is no guarantee of an exact correspondence, and a more precise interpretation of our procedure is that it identifies a low-dimensional subspace of sentence responses that successfully generalizes across participants. On this interpretation, sentence properties are best viewed as explaining different possible directions within this subspace. Indeed, if we allow sentence properties to have different directions within the two-dimensional space spanned by the PCs, we can achieve even higher correlations between sentence properties and the PCs (data not shown).

### 5. Modeling voxels using Sentence PCs

We sought to model voxel responses as a weighted sum of derived Sentence PCs. In particular, we were interested in conducting this modeling in a way that identifies representational content that is shared across participants.

We designed a leave-one-participant-out analysis in which we estimate Sentence PCs from all participants except one (seven participants) and use these PCs to predict voxel responses from the held-out participant. We used ordinary least squares (OLS) to determine optimal weights for characterizing voxel responses in the held-out participant:

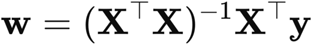

where **X** is a regressor matrix consisting of Sentence PCs (*n* sentence items by *m* Sentence PCs) as well as a constant term, **y** is an *n*-length column vector containing the held-out participant’s voxel’s response to each item, and **w** is an *m*-length column vector with the weights learned for each regressor.

We systematically varied the number of Sentence PCs included as regressors in the OLS model. To obtain unbiased estimates of the performance of the OLS model for each number of Sentence PCs, we used a cross-validation procedure where a subset of sentence items is used to estimate the weights **w** and the remaining sentence items are used to quantify model performance. We used a *k-*fold cross-validation scheme (with *k* = 5, results are quantitatively similar for *k* = 2 and *k* = 8). For *k* = 5, we fitted regression weights using 160 random sentence items (train set) and predicted the remaining 40 items (test set). We applied *z*-scoring to the regressors before fitting; regressors were *z*-scored using the mean and standard deviation identified on the train set, and the same transformation was applied on the test set. This procedure ensured independence (no data leakage) between the train and test sets. In each cross-validation fold, we computed the coefficient of determination (*R*^2^) between the predicted and measured voxel responses:

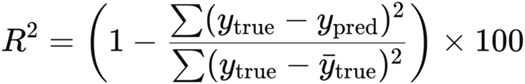

where *y_true_* and *y_pred_* are the measured and predicted voxel responses, respectively. The final *R*^2^value for each voxel was computed as the average across the *k* folds. Similarly, the fitted OLS weights were also averaged across the *k* folds. The weights indicate the contribution of each Sentence PC to a voxel’s response, and are analyzed in Figures 4 and 5. To meaningfully analyze Sentence PCs across participants, we checked that the Sentence PC signs were consistent across the eight held-out participant cross-validation iterations, which justifies aggregating across the weights learned for the eight participants. Note that one advantage of OLS procedure is that it does not assume anatomical correspondence across individuals, in contrast to some intersubject correlation approaches which have been used to study shared neural responses (e.g., Nastase et al. 2019).

To assess the significance of voxel predictions from the OLS prediction procedure, we repeated the OLS procedure but with the values within each Sentence PC permuted, in order to assess the null hypothesis that there is no relationship between Sentence PCs and brain responses. From the distribution of permuted *R*^2^ values (<u>SI 6</u>), we obtained the *R*^2^ value at a one-tailed *p*<.01 threshold (*R*^2^ = 1.06%) and all voxels predicted below this threshold were not analyzed.

Note that in the special case where Sentence PCs are derived using the naive approach for estimating the voxel covariance matrix (<u>Methods; Deriving Sentence PCs</u>; second approach for computing voxel covariance) and where the voxels to be modeled are the same as those from which the Sentence PCs are derived, it can be shown that the OLS solution for the weights are given simply by **w** = **V^T^**. Since we sought to investigate generalization of Sentence PCs to novel participants, we performed the explicit OLS fitting model procedure as described above.

### 6. Sentence properties

We collected a set of 16 sentence properties to characterize the identified Sentence PCs. Two properties were collected using behavioral experiments specifically for the current study (word-by-word reading times and concreteness of sentence meanings), while the remaining properties were obtained from behavioral experiments from prior work (Tuckute et al. 2024d) or extracted from text corpora.

First, to obtain a measure of how effortful each sentence is to process, we measured word-by-word reading times using the Maze task (Forster et al. 2009; Boyce et al. 2020). Second, building on the body of evidence for surprisal modulating language processing difficulty, in both behavioral psycholinguistic work (e.g., Demberg & Keller, 2008; Smith & Levy, 2013; Brothers & Kuperberg, 2021) and brain imaging investigations (e.g., Henderson et al. 2016; Willems et al. 2016; Heilbron et al. 2022; Shain, Blank et al. 2020, Shain et al. 2024b; Tuckute et al. 2024d), we computed the average surprisal for each sentence using three different models (surprisal is negative log probability; see the <u>Surprisal</u> section below). Third, motivated by the sensitivity of the language network to pointwise mutual information (PMI; Mollica, Siegelman et al. 2020), we computed the average PMI for each sentence. And fourth, we obtained 11 behavioral rating norms: 10 norms collected by Tuckute et al. (2024d), and one—for sentence concreteness—collected as part of this study. These behavioral norms spanned judgments related to both sentence well-formedness and content, and were grounded in prior work in psycholinguistics and neuroscience (Paivio et al. 1968; Kuchinke et al. 2005; Binder et al. 2005; Demberg & Keller 2008; Jack et al. 2013; Huth et al. 2016; Arfé et al. 2022).

#### Word-by-word reading times (Maze task)

The Maze task (Forster et al. 2009) is a two-alternative forced-choice task used to measure reading time, where participants choose between the correct word in a given sentence and a less plausible distractor word. Slower response times indicate words that are more effortful and time-consuming to process, and is used as a behavioral measure of online processing effort (Forster et al. 2009; Boyce & Levy 2023; Hao et al. 2023). We used a variant of the Maze task known as A(uto)-Maze, introduced by Boyce et al. (2020), which automatically generates distractor words using language models’ estimate of contextual surprisal. We implemented a more state-of-the-art language model (GPT2; Radford et al. 2018) compared to Boyce et al. (2020) to generate well-matched context-sensitive distractors (for related approach, see Heuser & Gibson (2023)). We used this A-Maze task to measure word-by-word reading times and computed the average reading time per sentence (as our fMRI data do not provide word-level responses).

#### Maze auto-generation process: Distractor word selection

We adopted the A-maze procedure introduced by Boyce et al. (2020) to identify distractors for the Maze task. For the surprisal computation, we used the pre-trained GPT2 from the HuggingFace library (Wolf et al. 2020; transformers version 4.38.2). For each position in the sentence besides the very first word, we computed the surprisal (in bits) of the correct next word (‘target’ word) and a candidate distractor word, given the preceding context. If a word was split into multiple sub-word tokens by the GPT2 tokenizer, we summed the log probabilities to obtain a single surprisal estimate per word. To identify distractor words that were poor continuations of the sentence, we computed the difference in surprisal between the target word and each candidate distractor. We ran A-Maze using the default surprisal parameters introduced in Boyce et al. (2020) (*min_delta* = 10 and *min_abs* = 25, where *min_delta* refers to how much more surprising the distractor should be than the target word in bits and *min_abs* refers to how surprising the distractors should be in bits) and evaluated 300 candidate distractor words for each position.

Following the A-Maze procedure (Boyce et al. 2020), and given that reading times are influenced by lexical frequency and word length (Kliegl et al. 2004), we matched distractor words to target words based on both frequency and length. For length matching, words with lengths between 4 and 10, the length was exactly matched. For words of length *n* < 4, distractor words, *x*, of length *n* <= *x* <= 4 were considered. For words of length *n* > 11, distractor words of length 11 <= *x* <= *n* were considered. Frequency matching was performed by first selecting candidate distractor words of the same log lexical frequency as the target word. If fewer than 300 words (number of words to evaluate for each word position) met this criterion, the frequency range was iteratively expanded to include words with frequencies ± *delta* from the original target frequency, where *delta* = 1, 2, 3, … until the number of candidate words that satisfied the criterion was greater than 300.

Following Boyce et al. (2020), the vocabulary for distractor selection was based on the English dictionary defined by the *wordfreq* Python package (version 3.1.1). In addition, to ensure well-formed distractor words, we further restricted the vocabulary to the top 30,000 most frequent English words in the SUBTLEXus corpus (Brysbaert et al. 2012), resulting in a final vocabulary of 24,778 words to sample distractors from.

Through manual inspection of the final distractor words, 36/1000 (3.6%) distractors generated from the algorithm were judged as poor distractors due to strong ambiguity. These 36 words were manually replaced using a word chosen from the set of 300 length- and frequency matched candidate distractors generated by the algorithm. In the final set of distractors, the distractor word length was exactly matched to the target word in 66.20% of word-distractor pairs, and within a difference of up to three characters for short words. 50.60% of the final selected distractors had log lexical frequencies within ± 2 of the target word (see <u>SI 13</u> for word length and frequency matching distributions). Finally, the difference in contextual surprisal between the target word and the distractor was 13.72 bits, on average (<u>SI 13</u>), indicating that the distractor generation procedure successfully identified distractors with higher contextual surprisal, as required for the Maze paradigm.

##### Experimental procedure and participants

The experiment was run using the Ibex-with-Maze implementation introduced in Boyce et al. (2020) and hosted on the Prolific crowd-sourcing platform. 40 participants were recruited. The study was restricted to individuals living in the USA who indicated English as their first and primary language.

We divided the 200 sentences into two random sets of 100 sentences each. Each participant was randomly assigned to either set one or set two. In total, a group of 20 participants were shown set one and a non-overlapping group of 20 participants were shown set two. Participants first provided informed consent, and then answered several demographic questions (whether English is their first language, which country they are from, and what age bracket they fall into); they were explicitly told that payment was not contingent on their answers to these questions. Participants were next presented with the survey-specific instructions and the following warning: “There are some sentences for which we expect everyone to answer in a particular way. If you do not speak English or do not understand the instructions, please do not do this hit—you will not get paid.”. Participants completed three practice items before proceeding with the main experiment.

Following the A-Maze paradigm, in each trial, participants were tasked with selecting between a word and a distractor for each word in the sentence. The first word was always paired with a nonce (x-x-x) distractor. For each subsequent word, the distractor was selected following the automatic procedure described above. In the case where participants made an incorrect choice, an error message was displayed, and participants were allowed to select the correct choice after a 1 second delay interval (to avoid terminating the sentence trial on an incorrect response; this variant of the Maze task was introduced as ‘error-correction Maze’ in Boyce & Levy (2023)).

The experiments were conducted with approval from and in accordance with MIT’s Committee on the Use of Humans as Experimental Subjects (protocol number 2010000243). The participants gave informed consent before starting each experiment and were compensated for their time (minimum US$12 per hour).

##### Participant and item exclusion

We collected a total of 24,000 word-distractor pair responses (40 participants × 100 sentences per participant × six words per sentence). For each word-distractor pair, as in Boyce et al. 2020, we recorded both the time to the first response, as well as the time to the correct response. For the remainder of this study, we refer to reading time (RT) as the time to the correct response, always excluding the 1 second delay period (unlike Boyce & Levy (2023), we do not exclude error and post-error RTs to maximize usable data from the short six-word-long sentences).

RT responses were excluded based on the following three pre-defined criteria: the first two from Boyce et al. (2020) and the last based on the other behavioral experiments in this study (<u>Methods; Behavioral norms</u>). First, we excluded the first word of each sentence since the first word was always paired with a nonce (x-x-x) distractor. This resulted in 20,000 word-distractor pair responses. Second, for each participant, we excluded sentences with RT outliers: specifically, if any word had a first time to response < 100 ms (likely caused by unintentional selections prior to reading), or if the time to either the first or correct response exceeded 5,000 ms (likely due to distraction). This procedure left 19,440 word-distractor pair responses. Third, we excluded participants due to low correlations with the remaining participants (same criterion as the other behavioral experiments). Specifically, participants whose average correlation with all other participants was more than two standard deviations below the average were excluded. The inter-participant correlation was computed using the vector of average word-level RTs per sentence. Two participants were excluded, leaving a total of 38 participants and 18,520 word-distractor pair responses for subsequent analyses. Finally, to obtain the average RT for each of the 200 sentences, we averaged the word-distractor pair RTs within a given sentence across participants.

The experiment took 25.37 minutes, on average (SD across participants = 15.40). Each item was rated by 18.52 participants (SD across items = 0.80). The average inter-participant Pearson correlation, computed by correlating a vector of average RTs per sentence for a given participant with the vector of average sentence-level RTs for each of the remaining participants and taking the average of these pairwise correlation values, was 0.30 (SD across participants = 0.06) for set one and 0.40 (SD = 0.06) for set two. We also computed inter-participant correlations using the word-level RTs, by correlating a vector of word-distractor pair responses for a given participant with the vector of responses for each of the remaining participants and taking the average of these pairwise correlation values. The word-level correlation was 0.31 (SD across participants = 0.03) for set one and 0.38 (SD = 0.05) for set two. Of the 38 participants, 36 participants had an accuracy >90%, where accuracy is defined as the proportion of times a participant selected the proper word given the linguistic context over the distractor word. The remaining two participants had accuracies of 88.4% and 89.0%.

#### Surprisal

We estimated the negative log probability of each word given its context within a sentence. The negative log probability of a word/sentence is known as “surprisal” (Hale 2001). The surprisal of each sentence was computed using three surprisal models aimed at capturing distinct information about the predictability of each sentence, as detailed below:

##### Large language model (LLM) surprisal

LLM surprisal was computed using GPT2-XL, a unidirectional-attention language model trained to predict the upcoming word by attending to prior words in the sequence through the attention mechanism (Vaswani et al. 2017). This training procedure makes the surprisal estimates sensitive to various aspects of all the preceding words in the sentence (Radford et al. 2018). We used the pre-trained GPT2-XL from the HuggingFace library (Wolf et al. 2020; transformers version 4.11.3). GPT2-XL was trained on 40GB on web text from various domains (WebText dataset). Each sentence was tokenized using the model’s standard tokenizer (GPT2Tokenizer) and the special token, [EOS], was prepended to each sentence. Punctuation was retained. Surprisal was computed in natural log units (nats). We obtained the sentence-level surprisal by taking the mean of the token-level surprisals.

#### Lexical n-gram (5-gram) surprisal

Lexical surprisal was computed using a 5-gram language model which estimates surprisal given the preceding four words (i.e., for the first four words there is no such window; the surprisal for the first word of a sentence is the probability of that given word beginning the sentence, for the second word it is the 2-gram probability and so on). The model was trained using the KenLM library (Heafield, 2011; build on 12/24/2022; https://github.com/kpu/kenlm) with default smoothing parameters (modified Kneser-Ney smoothing) on the training set of Wikitext-103 (Merity et al. 2016). The training data were tokenized using the sent_tokenizer from the NLTK library (Bird & Loper, 2004; version 3.8.1) followed by tokenization of sentences using the wordpunct_tokenizer from the same library. Punctuation was stripped and all words were lower-cased. Surprisal estimates for each word in the materials was obtained using default parameters of the full_scores function in KenLM. Surprisal was computed in base-10 log units. Surprisal for each word in the sentence along with an end-of-sentence token (aaa/sbbb) was computed. 6 unique words from our materials (out of a total of 670 unique words, 0.9%) were out of vocabulary for the n-gram model and hence surprisal could not be estimated for these words. We obtained the sentence-level surprisal by taking the mean of the word-level surprisals.

##### Lexicalized probabilistic context-free grammar (PCFG) surprisal

PCFG surprisal was computed using the incremental left-corner parser of van Schijndel et al. (2013) trained on a generalized categorial grammar reannotation (Nguyen et al. 2012) of Wall Street Journal sections 2 through 21 of the Penn Treebank. Each sentence was tokenized using a Penn Treebank Tokenizer and punctuation was retained. Surprisal was computed in base-2 log units (bits). We obtained the sentence-level surprisal by taking the mean of the word-level surprisals.

#### Pointwise mutual information (PMI)

PMI was computed to quantify the association among nearby words based on co-occurrence counts, a metric from information theory (Church & Hanks 1990; Jurafsky & Martin 2008). We utilized the Corpus of Contemporary American English (COCA; Davies, 2010). Punctuation was stripped and all words were lower-cased. We used a six-word sliding window to obtain the co-occurrence counts among pairs of words. We selected the top 20,000 most frequent words to form our vocabulary based on individual word frequencies. PMI was computed using the formula:

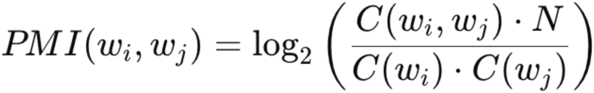

where *C(w_i_, w_j_)* is the co-occurrence count of words *w_i_* and *w_j_, C(w_i_)* and *C(w_j_)* are their individual counts, and *N* is the total number of words in the corpus. Negative PMI values were set to zero to obtain positive PMI because negative PMI values are in practice extremely noisy due to data sparsity (Jurafsky & Martin 2008). We obtained the sentence-level PMI by taking the mean of the PMI values for all unique word pairs within a sentence.

#### Behavioral norms

We obtained behavioral rating norms for each sentence (11 in total). Ten of these norms were collected by Tuckute et al. (2024d) and the final norm—sentence concreteness—was collected as part of the current study.

##### Ratings from prior work (10 sentence properties)

We used the ratings for 10 sentence properties collected by Tuckute et al. (2024d): plausibility, grammaticality, general frequency, conversational frequency, places, physical objects, mental states, imageability, arousal, and valence. These ratings were collected for a total of 2,000 sentences, of which our 200 sentences were a subset. We refer to the methods in Tuckute et al. (2024d) for detailed information. In brief, participants from crowd-sourced platforms were asked to rate each sentence according to a given property (e.g., *grammaticality*: “How much does the sentence obey the rules of English grammar?” or *imageability*: “How visualizable is the sentence’s content?”) on a Likert scale from 1 to 7.

##### Rating collected for the current study (1 sentence property)

We collected a rating targeted at assessing the concreteness of a sentence’s meaning. Participants were asked to rate each sentence according to “How much is the sentence’s meaning tied to perceptual experience?” on a Likert scale from 1 to 7 (see full instructions in <u>SI 14A)</u>. We followed the same procedure and pre-defined exclusion criteria as the ratings collected by Tuckute et al. (2024d), as detailed below.

Participants were recruited using the Prolific crowd-sourcing platform using the same inclusion criteria as in the Maze task (<u>Methods; Word-by-word reading times</u>). As in the Maze task, participants provided informed consent, completed the same demographic questions, and received the task instructions (<u>SI 14A</u>).

40 participants took part in the experiment. Four participants were excluded following pre-defined exclusion criteria (<u>SI 14B</u>), leaving 36 participants (90%). In particular, four participants were excluded based on a low correlation with the remaining participants. The experiment took 13.85 minutes, on average (SD across participants = 0.66). Each item was rated by 18 participants (SD across items = 0.00). The average inter-participant Pearson correlation, computed by correlating a vector of responses for a given participant with the vector of responses for each of the remaining participants and taking the average of these pairwise correlation values, was 0.38 (SD across participants = 0.11). The experiments were conducted with approval from and in accordance with MIT’s Committee on the Use of Humans as Experimental Subjects (protocol number 2010000243). The participants gave informed consent before starting each experiment and were compensated for their time (minimum US$12 per hour).

