## Supplemental Information for "A two-dimensional space of linguistic representations shared across individuals"

#### SI 1: Sentence PCs derived using different voxel inclusion criteria and different decomposition parameters

The Sentence PCs are derived through matrix decomposition of a data matrix consisting of beta weights in percent signal change. There are three main decision points in this procedure: (1) Which voxels are included, (2) How the voxel covariance is computed, and (3) The decomposition method.

As described in the main text (Methods; Deriving Sentence PCs), we included voxels with a noise ceiling signal-to-noise ratio (NCSNR) above 0.4 within the language parcels and further randomly subsampled 75% of these voxels. For computing the voxel covariance matrix, we use Generative Modeling of Signal and Noise (GSN; Kay et al., 2024) to isolate the covariance of the signal, excluding noise defined as variability across trial repetitions (the alternative is to naively compute the covariance of the data, which is affected by both signal and noise). We then applied singular value decomposition (SVD) to the resulting signal covariance matrix. These choices are summarized below:

1) **Voxel inclusion:** NCSNR > 0.4 in language parcels; 75% subsampling. **Covariance method:** Signal covariance matrix. **Decomposition method:** SVD.

In these supplementary analyses, we vary the three key choices of the main analysis: voxel inclusion, covariance method (signal covariance matrix or naive covariance matrix), and decomposition approach (SVD or non-negative matrix factorization (NMF)). Data from all eight participants were used to derive Sentence PCs in these analyses. We examine the following six additional configurations:

2) **Voxel inclusion:** NCSNR > 0.4 in language parcels; 75% subsampling. **Covariance method:** Naive covariance matrix. **Decomposition method:** SVD.
3) **Voxel inclusion:** NCSNR > 0.3 in language parcels; 40% subsampling. **Covariance method:** Signal covariance matrix. **Decomposition method:** SVD.
4) **Voxel inclusion:** NCSNR > 0.3 in language parcels; 50% subsampling. **Covariance method:** Naive covariance matrix. **Decomposition method:** SVD.
5) **Voxel inclusion:** Select top 10% NCSNR voxels in language parcels for each participant; 80% subsampling. **Covariance method:** Signal covariance matrix. **Decomposition method:** SVD.
6) **Voxel inclusion:** Select voxels with a *sentences* > *nonwords* *t*-statistic of at least 3 in either a visual or auditory language localizer experiment in language parcels; 40% subsampling. **Covariance method:** Signal covariance matrix. **Decomposition method:** SVD.
7) **Voxel inclusion:** NCSNR > 0.4 in language parcels; 50% subsampling. **Covariance method:** Naive covariance matrix. **Decomposition method:** NMF (see NMF methods in the last part of this supplemental information).

The heatmap below shows the correlation of the first 10 Sentence PCs obtained across all seven configurations (their numbers correspond to the rows/columns in the figure; note that the PCs here are zero-indexed). Critically, the first few Sentence PCs remain highly robust to the modified parameters (voxel inclusion, covariance method, and decomposition method). Two minor points are worth noting: (1) There is a sign flip for Sentence PC 2 for configuration 6, resulting in a strong negative correlation with Sentence PC 2 in configuration 1 ($r$ = -0.87) instead of positive correlation—this is not a problem since the sign of SVD-derived components is arbitrary, (2) NMF yields components that are sorted differently, but Sentence PCs strongly correlated with the first and second PCs from configuration 1 are still present.

**A** Correlation of Sentence PCs obtained from different voxel inclusion criteria, covariance computations, and decomposition methods

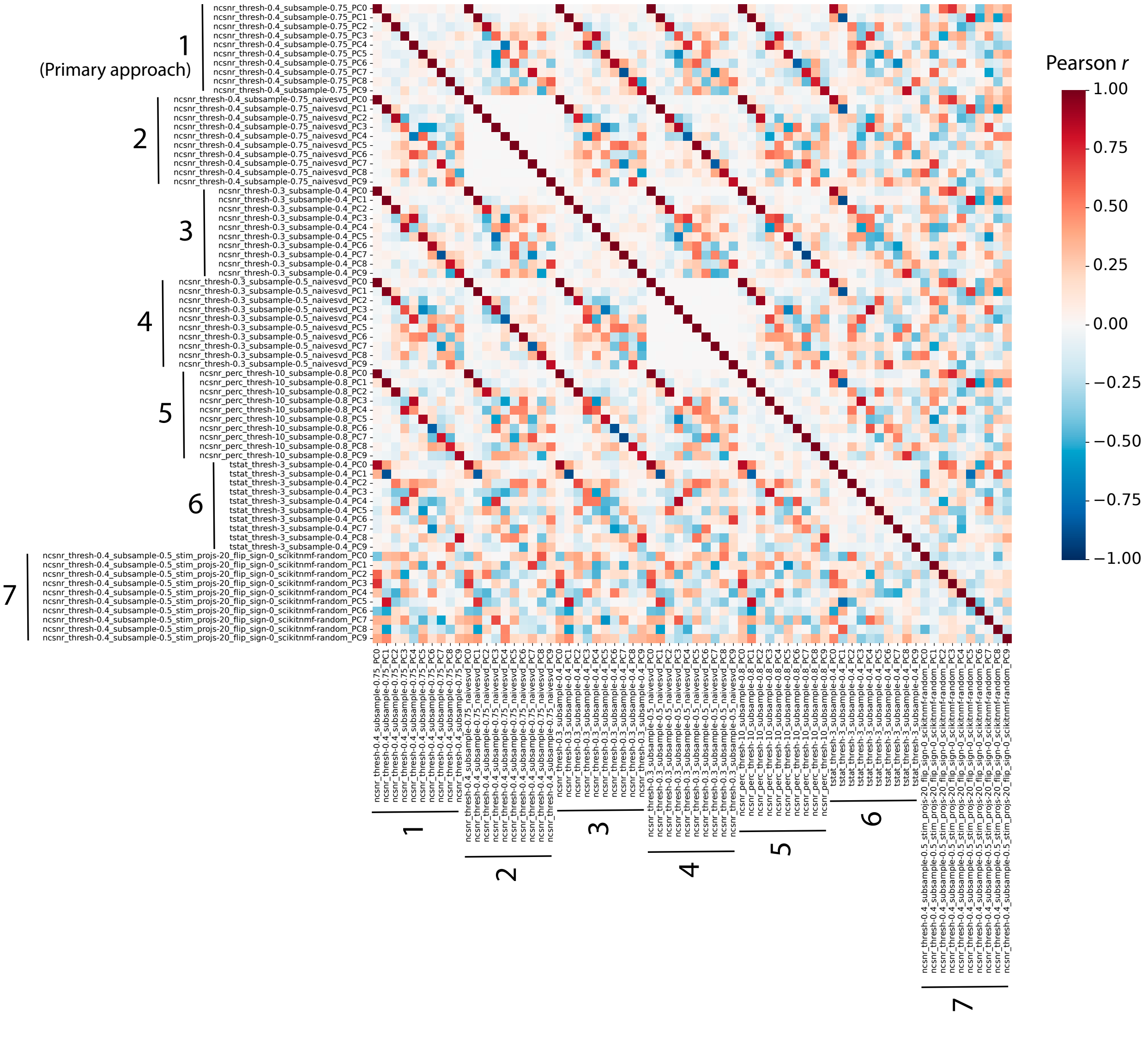

*Non-negative matrix factorization (NMF) procedure*

Here, we provide the methods used to derive the NMF components used in configuration 7. NMF was implemented in scikit-learn (version 1.3.2; Pedregosa et al., 2011) with 'random' initialization and a maximum of 1,000 iterations.

First, the data were preprocessed: Similar to Khosla et al. (2022), we preprocessed the data matrix to make it suitable for NMF by performing a baseline shift of voxel responses by subtracting the minimum z-scored response of each voxel (across all sentence items) from its responses to all items.

Second, we determined the optimal number of NMF components by performing NMF decomposition with 1 to 20 components. For each number of components, we conducted 10 independent NMF "stability" runs with different random initializations to account for stochasticity in the algorithm. We calculated the Bayesian Information Criterion (BIC) for each number of components and averaged these values across the stability runs. The number of components with the lowest average BIC was selected as the optimal number of components (13 components were identified from $n$ = 8 participants).

Third, we ensured that we only obtained stable NMF components. We aggregated the component matrices from all stability runs corresponding to the optimal number of components. We used the "IsolationForest" method from scikit-learn with default parameters to retain the stable components. Then, we performed $k$-means clustering on the filtered components, setting the number of clusters equal to the optimal number of components. Within each cluster, we averaged the components to obtain representative components. Finally, we sorted these components based on cluster size, assuming that clusters with more components represent more stable and significant patterns in the data (where NMF components correspond to "Sentence PCs" in the main analyses).

#### SI 2: Predictivity of Sentence PCs for different voxel inclusion criteria

**A** Predictivity ($R^2$) of Sentence PCs across different voxel inclusion criteria

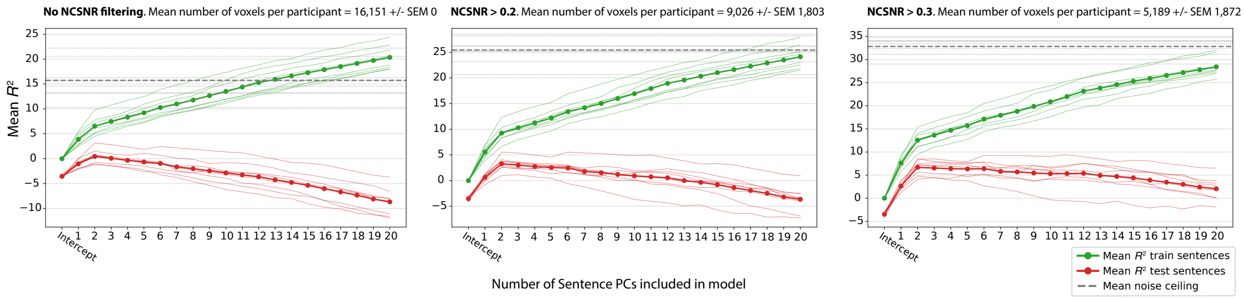

Figure 2B in the main text shows the predictivity (coefficient of determination, $R^2$) of Sentence PCs for voxels in the language parcels with NCSNR (noise ceiling signal-to-noise ratio) > 0.4. These plots show the predictivity in the language parcels with no NCSNR filtering, NCSNR > 0.2, and NCSNR > 0.3. Critically, the finding that two components are optimal remains robust regardless of voxel inclusion criteria. The x-axis shows the number of Sentence PCs included in the OLS model, starting with 0 (intercept only). Red lines show the $R^2$ test performance, green lines show the $R^2$ train performance. Thick lines show the average $R^2$ across $n$ = 8 participants, thin lines show $R^2$ for individual participants. The horizontal gray lines show the noise ceiling: the thick line shows the average noise ceiling across participants, thin lines show noise ceilings for individual participants.

#### SI 3: Comparison of model predictivity and language selectivity

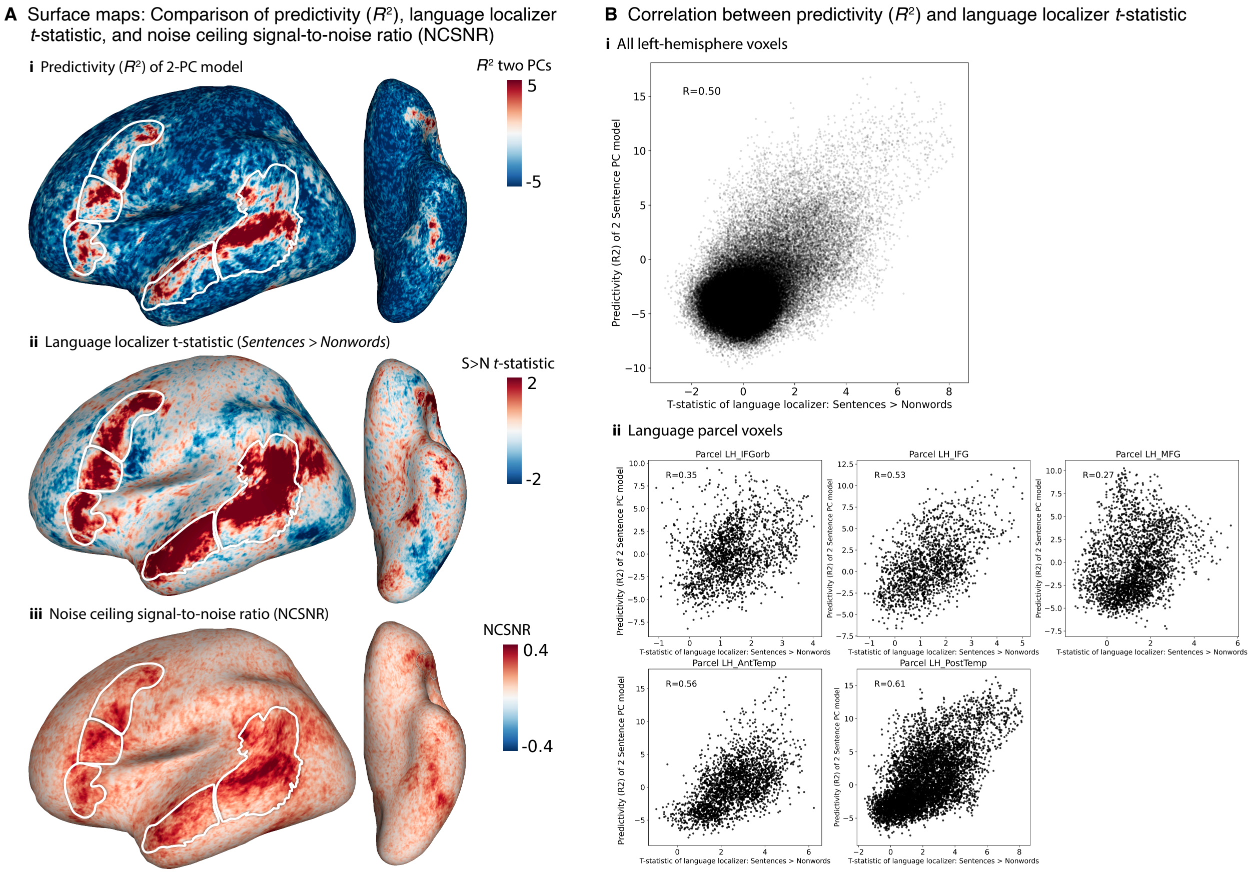

Panel **A** shows three surface maps on the left lateral and ventral surface in the fsaverage space (Fischl et al., 1999) each averaged across $n$ = 8 participants. Panel **i** shows the predictivity of the two Sentence PC model, panel **ii** shows the language selectivity, and panel **iii** shows the noise ceiling signal-to-noise ratio (NCSNR). The $R^2$ map (panel i) is the same as in Figure 2C in the main text. The language localizer $t$-statistic in panel **ii** was obtained from a well-validated localizer task (Fedorenko et al. 2010), contrasting meaningful language (sentences) with a perceptually matched control (lists of nonwords) (see Methods; Language network localizer experiment).

Panel **B** shows the relationship between language localizer $t$-statistic (x-axis) and model predictivity of the two Sentence PC model (y-axis) for all voxels in the left hemisphere (panel **i**) and for each of the five anatomical language parcels (panel **ii**; illustrated as the white outlines in panel **A**). The correlations and scatter plots were based on the average of $n$ = 8 participants. Generally, there is a moderate correlation between language selectivity and predictivity.

#### SI 4: Predictivity of Sentence PCs and weights onto Sentence PCs in the right hemisphere

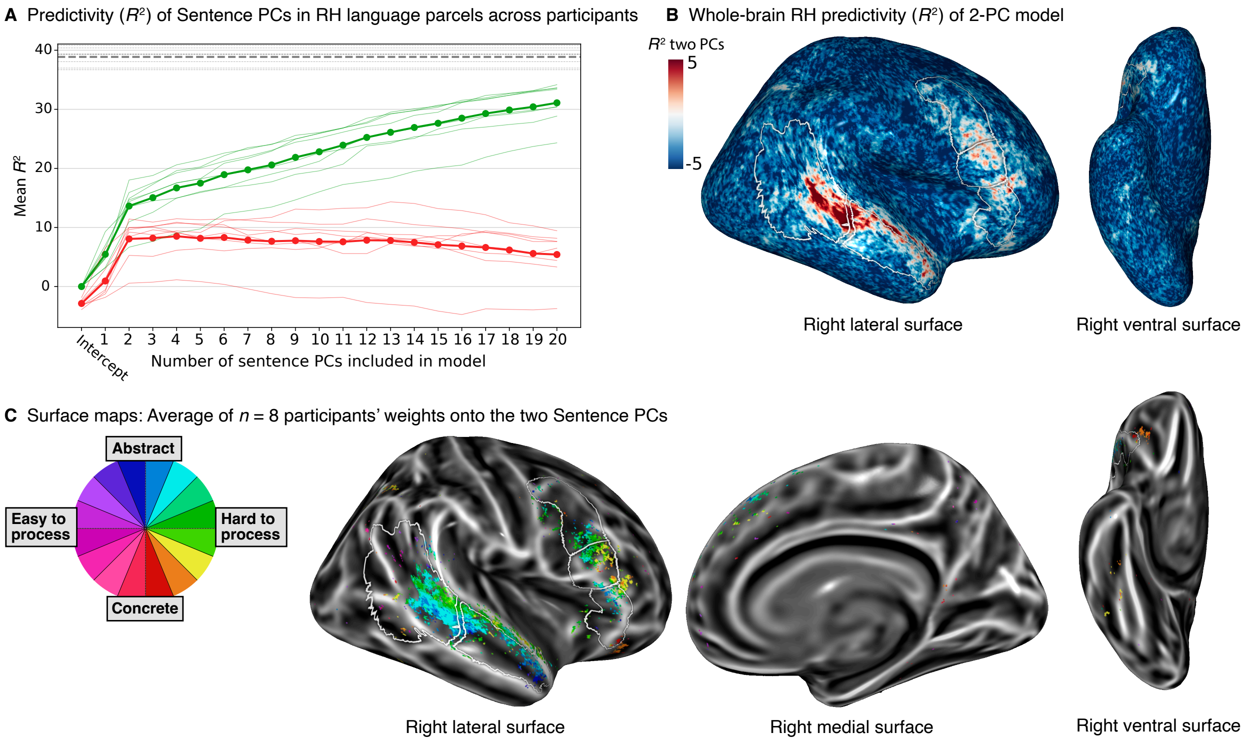

All analyses in the main text focus on the left hemisphere (LH), motivated by the left-lateralization of the language network (e.g., Lipkin et al., 2022). Here, we provide some key analyses of the right hemisphere (RH), using the Sentence PCs derived from the LH. Panel **A** shows the predictivity (coefficient of determination, $R^2$) of Sentence PCs for voxels in the language parcels with NCSNR (noise ceiling signal-to-noise ratio) > 0.4 (on average 1,061 voxels per participant +/- SEM 807). The mean $R^2$ across participants for two PCs was 8.07 +/- SEM 1.24. Note that the relative contribution of PC2 was numerically larger in the RH compared to the LH (Figure 2B main text), albeit not statistically significant ($t$=-1.77, $p$=0.12 via two-tailed paired $t$-test). Panel **B** shows the predictivity plotted on the inflated right hemisphere surface (in the fsaverage space; Fischl et al., 1999). The $R^2$ values were averaged for every voxel across $n$ = 8 participants. Panel **C** shows surface maps for the average of $n$ = 8 participants' weights onto the two Sentence PCs for significantly predicted voxels for at least three participants (mirroring Figure 4A in the main text). The "rainbow" colormap represents the two-dimensional space spanned by the two Sentence PCs, where the *direction* (angle) of a voxel in this space reflects the contribution of the two PCs in predicting that voxel's response. The spatial topography observed in the RH appears qualitatively similar to that in the LH (Figure 4A in the main text), with the more superior part of the temporal lobe preferring hard-to-process sentences (green colors) and the more medial and anterior part being more abstract-preferring. Overall, the predictive power of LH-derived Sentence PCs in the RH suggests that the same underlying representational dimensions are shared across hemispheres.

#### SI 5: Inter-correlation among sentence properties

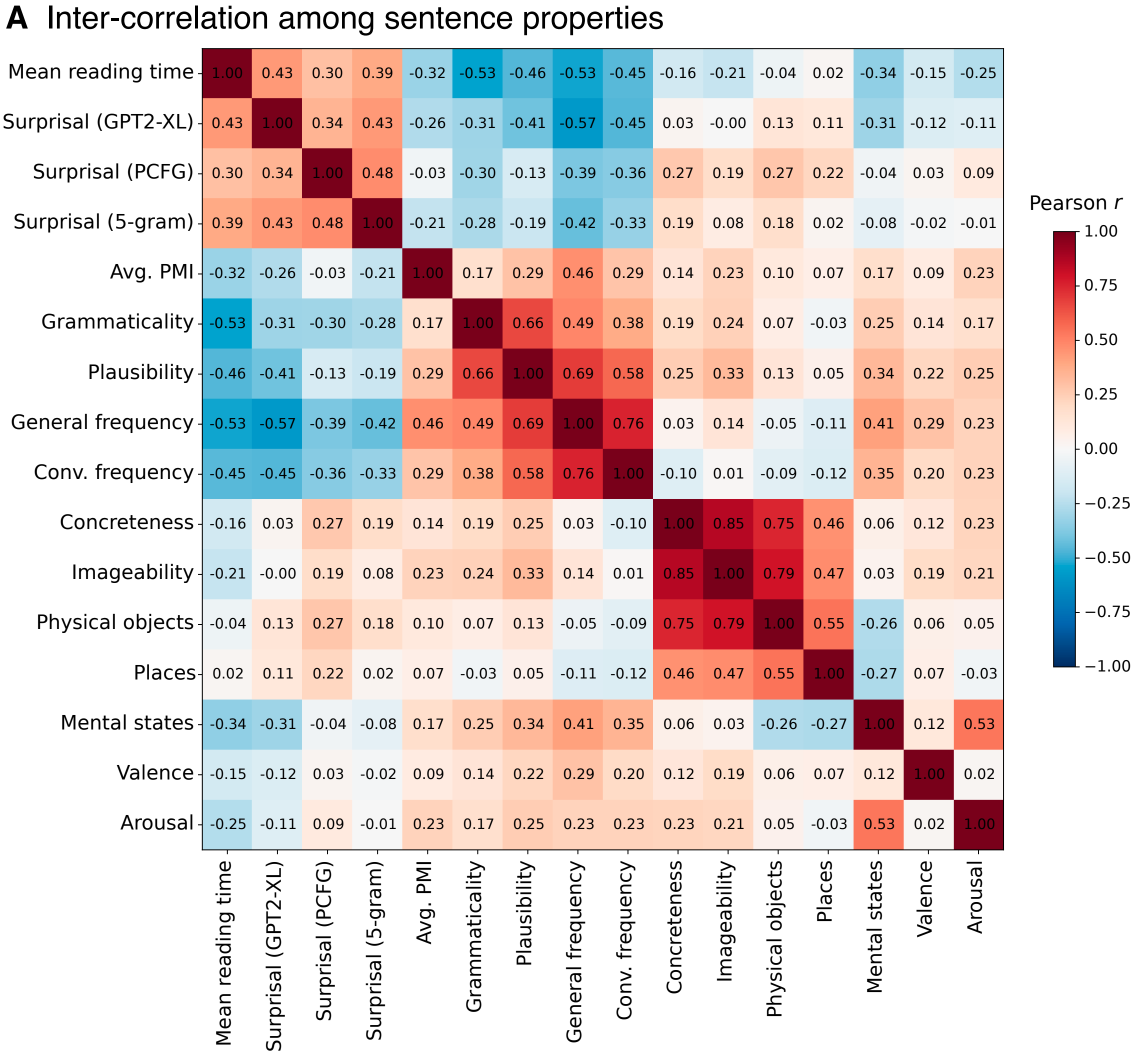

The heatmap shows the Pearson correlation among all the sentence properties (used in Figure 3 in the main text).

#### SI 6: Permuted predictivity of Sentence PCs

**A** Permuted predictivity ($R^2$) of Sentence PCs across participants

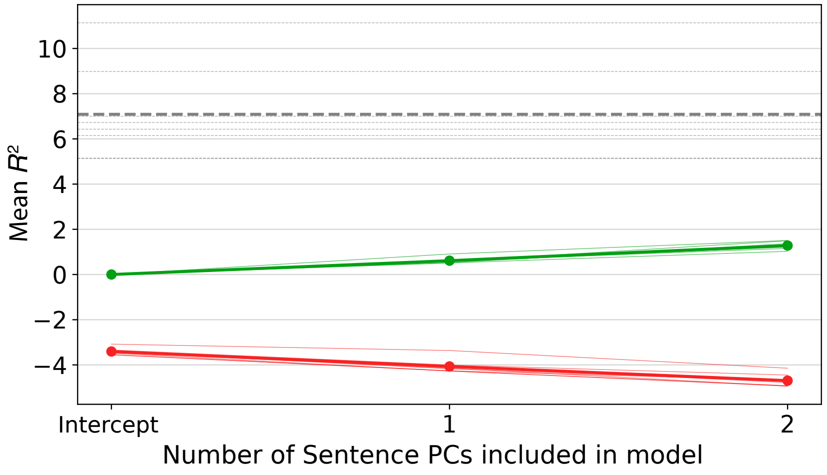

Predictivity (coefficient of determination, $R^2$) for OLS models with an increasing number of Sentence PCs used as regressors to predict voxels in a held-out participant. In contrast to Figure 2 in the main text, the values within each Sentence PC were permuted, in order to assess the null hypothesis that there is no relationship between Sentence PCs and brain responses. The x-axis represents the number of Sentence PCs included in the OLS model, starting with 0 (intercept only; note that because we were interested in the two Sentence PC model, we only iterated over up to two PCs as regressors). Red lines show the $R^2$ test performance, green lines show the $R^2$ train performance. Thick lines show the average $R^2$ across $n$ = 8 participants, thin lines show $R^2$ for individual participants. All voxels in the left hemisphere were included for each participant.
From the distribution of randomly permuted $R^2$ values across participants, we obtained the $R^2$ value that is significantly greater than chance at a one-tailed $p$<.01 threshold ($R^2$ = 1.06%).

#### SI 7: Reliability of voxel-level angle metric

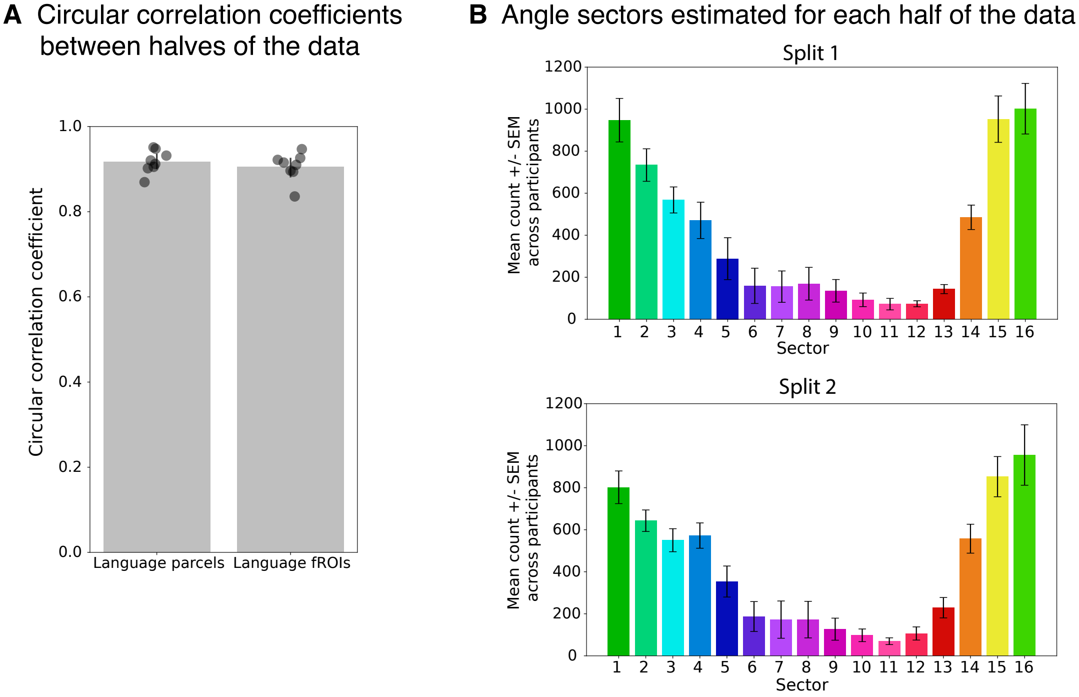

We evaluated the reliability of the voxel-level angle metric by examining the extent to which the OLS weights learned onto the two Sentence PCs are stable across stimuli. To do so, we randomly split the sentence items in half (100/100 sentences) for each participant. We learned the weights onto the two Sentence PCs for each split of the data. Then, we computed the circular correlation coefficient between the significantly predicted voxels (thresholded at $R^2 > 1.06\%$ using the average of the $R^2$s obtained from the two splits of the data) in either the language parcels or the language fROIs (panel **A**). The circular correlation coefficient was defined as (Jammalamadaka et al. 2001):

$$\rho = \frac{\sum \sin\left(\theta_{1_i} - \bar{\theta}_1\right)\ \sin\left(\theta_{2_i} - \bar{\theta}_2\right)}{\sqrt{\sum \sin^2\left(\theta_{1_i} - \bar{\theta}_1\right)\ \sum \sin^2\left(\theta_{2_i} - \bar{\theta}_2\right)}}$$

where $\theta_{1i}$ and $\theta_{2i}$ are the two sets of angular measurements (in radians) for the $i$th observation (voxel), and $\bar{\theta}_1$ and $\bar{\theta}_2$ are the circular means.

Panel **B** shows the quantification of angle sectors for voxels in the language parcels, as estimated on each half of the data. Error bars denote SEM across participants. Each histogram includes 51,591 voxels total (mean voxels per participant: 6,449 +/- 1,625 SEM). These analyses illustrate that the voxel-level angles and corresponding sectors were stable across two independent splits of stimuli.

#### SI 8: Surface maps: Participant-wise weights onto the two first Sentence PCs

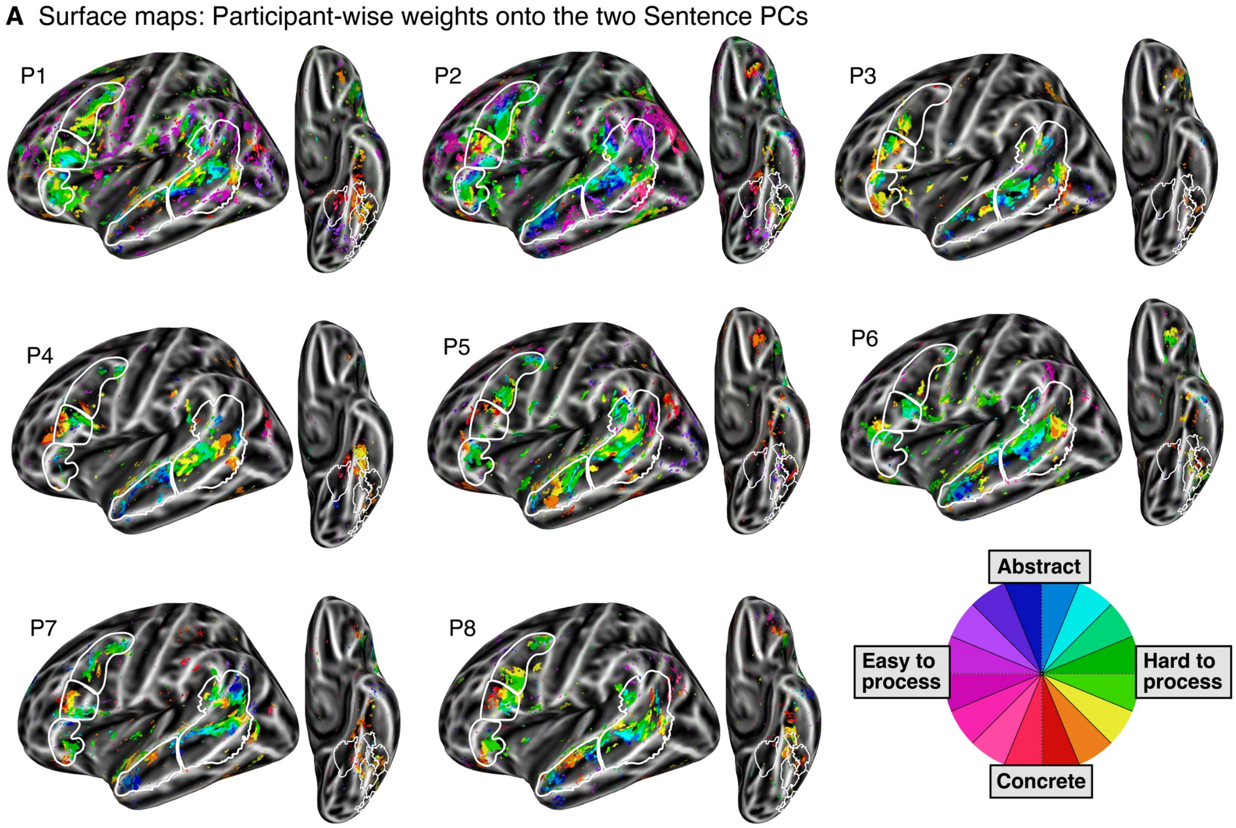

Surface maps for all eight individual participants. As in Figure 4 in the main text, the maps show the weights onto the two Sentence PCs using the "rainbow" sector color scheme. The "rainbow" colormap represents the two-dimensional space spanned by the two Sentence PCs, where the *direction* (angle) of a voxel in this space reflects the contribution of the two PCs in predicting that voxel's response. The white outlines on the surface map delineate the language parcels. Voxels that were significantly predicted are shown (same threshold as Figure 4, $R^2 > 1.06\%$).

#### SI 9: Participant-level consistency of angle metric

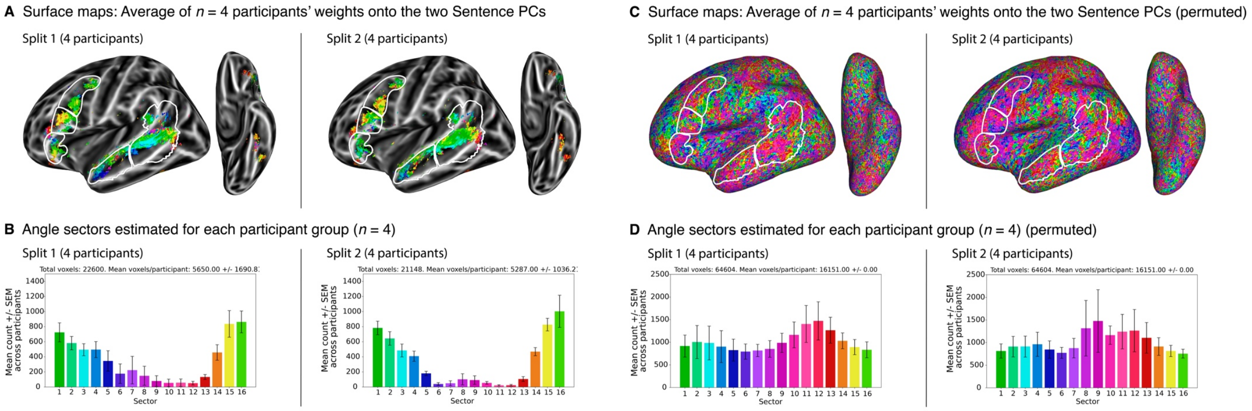

We evaluated the stability of the voxel-level angle metric across participants by examining the extent to which random subsets of participants exhibit similar angle tuning profiles. To do so, we randomly split the participants in half (4/4 participants) and visualized the average of $n$ = 4 participants' weights onto the two Sentence PCs on the brain surface (panel **A**), accompanied by the corresponding histogram that quantify the angle sector values in the anatomical language parcels (panel **B**). Both the surface maps and histograms use the same "rainbow" colormap as in the main text. The white outlines on the surface map delineate the language parcels. The surface maps include voxels with an average $R^2 > 1.06\%$ across all eight participants. The histograms include voxels that were significantly predicted within each individual, using the same $R^2 > 1.06\%$ threshold. These plots illustrate that the angle metric is consistent across independent groups of participants.

As a control for the angle tuning profiles shown in panels A and B, panels **C** and **D** present the exact same analyses using weights estimated from an OLS model in which values within each Sentence PC were permuted (see also SI 6). The permuted plots were not filtered by any $R^2$ threshold. As evident from the permuted plots, the patterns observed in panels A and B are no longer present and do not replicate across independent groups of four participants.

### SI 10: Quantification of Sentence PC weights in the left ventral surface

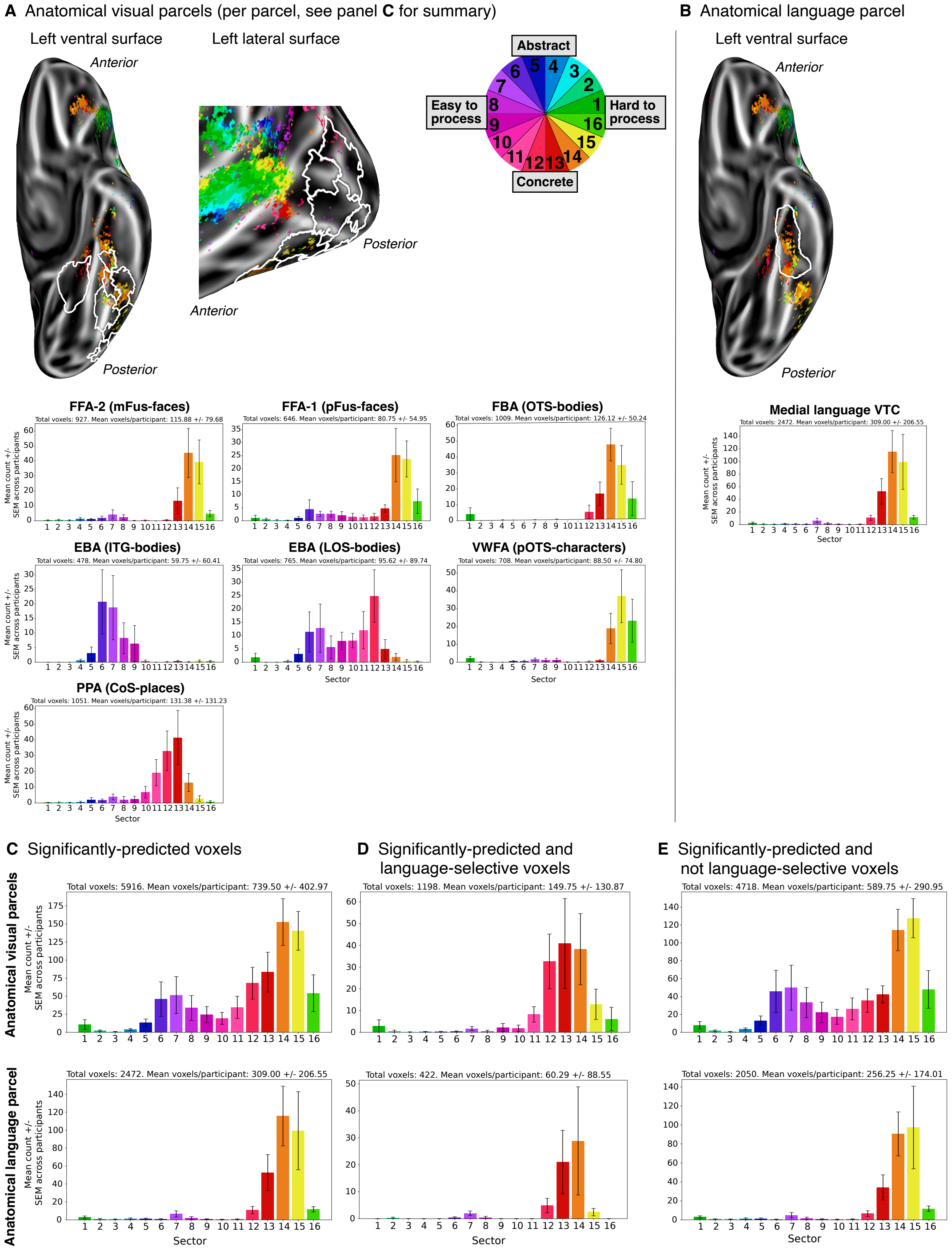

Panel **A** shows the quantification of angle sectors for voxels within each anatomical probabilistic visual parcel (Rosenke et al., 2021; see Methods; Visual parcels). The surface maps show the average of $n = 8$ participants' weights onto the two Sentence PCs for significantly predicted voxels ($R^2 > 1.06\%$). Identical to Figure 4 in the main text, the surface maps only show voxel locations where at least three participants exhibited a significantly predicted voxel above the significance threshold. The white outlines on the surfaces demarcate the parcel locations (10 in total). The histograms quantify the angle sectors in individual parcels. Three parcels are not shown because they exhibited fewer than 5% significantly predicted voxels (IOG-faces, IOS-characters, and hMT-motion), leaving seven parcels. Error bars in all histograms denote SEM across participants. Overall, visual parcels tend to prefer concrete sentences, though their tuning profiles vary.

Panel **B** shows the quantification of angle sectors for voxels within an anatomical probabilistic language parcel defined in the ventral temporal cortex (VTC) based on auditorily presented sentences and degraded speech (see Methods; Ventral language parcel; Li et al. 2024). This VTC language parcel partly overlapped with visual parcels FFA-2 and FBA. As in panel A, the surface map shows the average of $n = 8$ participants' weights onto the two Sentence PCs for significantly predicted voxels, with the white outline demarcating the parcel location. The histogram quantifies the angle sectors in the VTC language parcel. The VTC language parcels tends to prefer concrete and hard-to-process sentences.

Panels **C**, **D**, and **E** break down the significantly predicted voxels (panel **C**) into language-selective voxels (panel **D**) and non-language-selective voxels (panel **E**). Language-selectivity was defined as a *sentences* > *nonwords* $t$-statistic > 2 (see Methods; Language network localizer experiment); voxels with $t$-statistics < 2 were classified as non-language-selective. The first row shows the anatomical visual parcels, and the second row shows the VTC language parcel (note that in panel D, one participant did not have any significantly predicted and language-selective voxels in the language parcel, hence the plot only includes data from 7 participants). Both language-selective and not language-selective voxels generally show a preference for concrete sentences.

#### SI 11: Quantification of Sentence PC weights grouped into four sectors (hard-to-process, abstract, easy-to-process, and concrete)

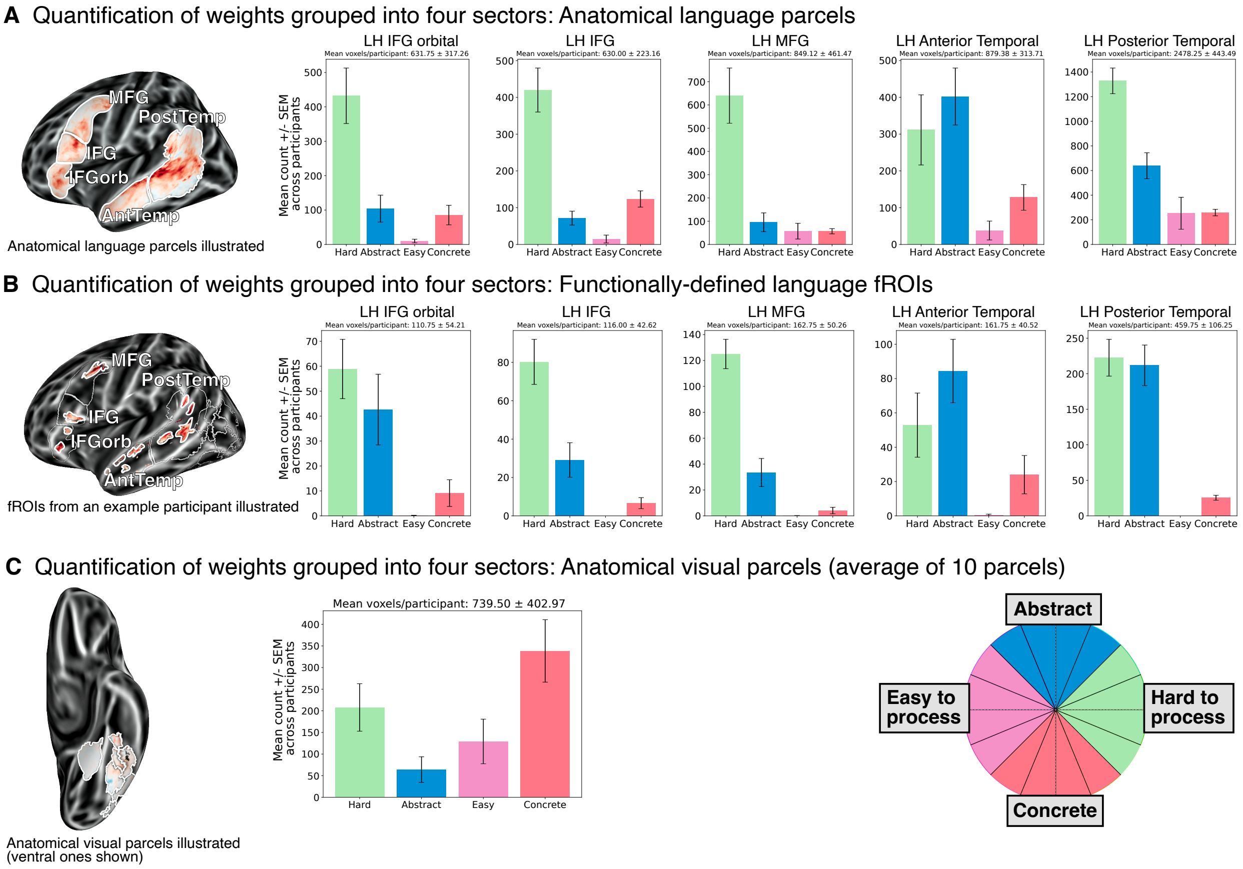

This figure summarizes the voxel tuning by grouping the 16 angle sectors into four broader categories: hard-to-process (green), abstract (blue), easy-to-process (pink), and concrete (red), as illustrated in the color wheel (bottom right). Panel **A** shows the quantification for voxels within anatomical language parcels, panel **B** for language fROIs, panel **C** for the average of the 10 anatomical probabilistic visual parcels. Panels A and B present the same data as in Figure 5 in the main text, while panel C presents the same data as Figure 4D in the main text. Error bars denote SEM across participants. All language parcels and fROIs generally prefer hard-to-process and abstract sentences, with the temporal parcels/fROIs showing a larger proportion of abstract-preferring voxels compared to the frontal areas. The language parcels/fROIs have a low proportion of voxels preferring concrete or easy-to-process sentences. In contrast, the probabilistic visual parcels show an overall preference for concrete sentences.

#### SI 12: Quantification of Sentence PC weights with the average angle per fROI

**A** Voxel-level angles in functionally-defined language fROIs with average angle

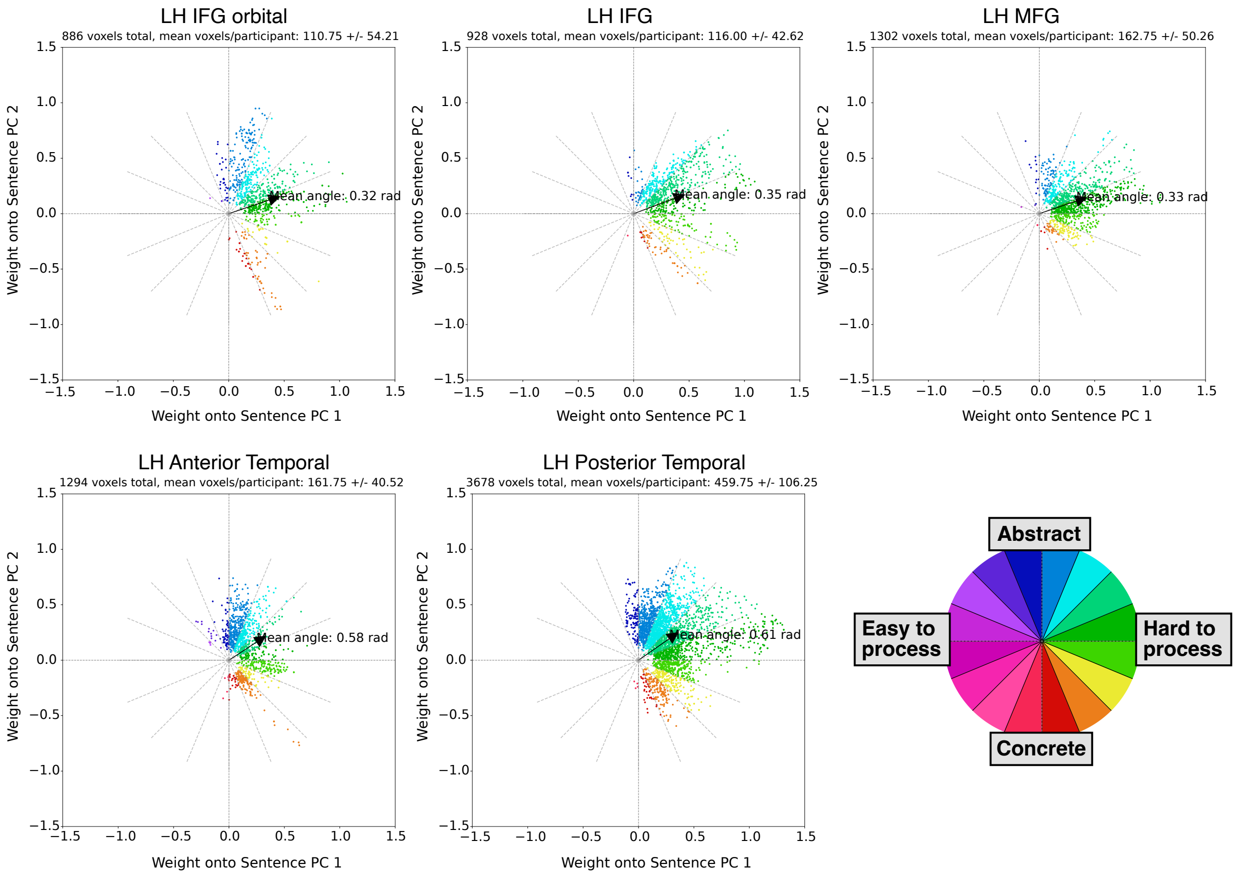

The figure shows the angle sectors for voxels within the five functionally-defined language fROIs (Fedorenko et al. 2010; see Methods; Language network fROIs) including the average angle per fROI (these scatter plots supplement the histograms in the main text, Figure 5B). The plots show the weights for each voxel onto Sentence PC 1 (x-axis) and Sentence PC 2 (y-axis) across $n = 8$ participants for significantly predicted voxels ($R^2 > 1.06\%$). The average angle per fROI was computed by first obtaining the average angle within each individual, and then averaging those angles to equalize participant contributions. It is evident that the mean angle is roughly similar across fROIs, but the internal *composition* of voxel tuning profiles varies among fROIs: for instance, the anterior temporal fROI contains a large proportion of voxels preferring "Abstract" sentences (dark blue colors) while IFG does not.

#### SI 13: Behavioral experiment: Quality of Maze distractor words

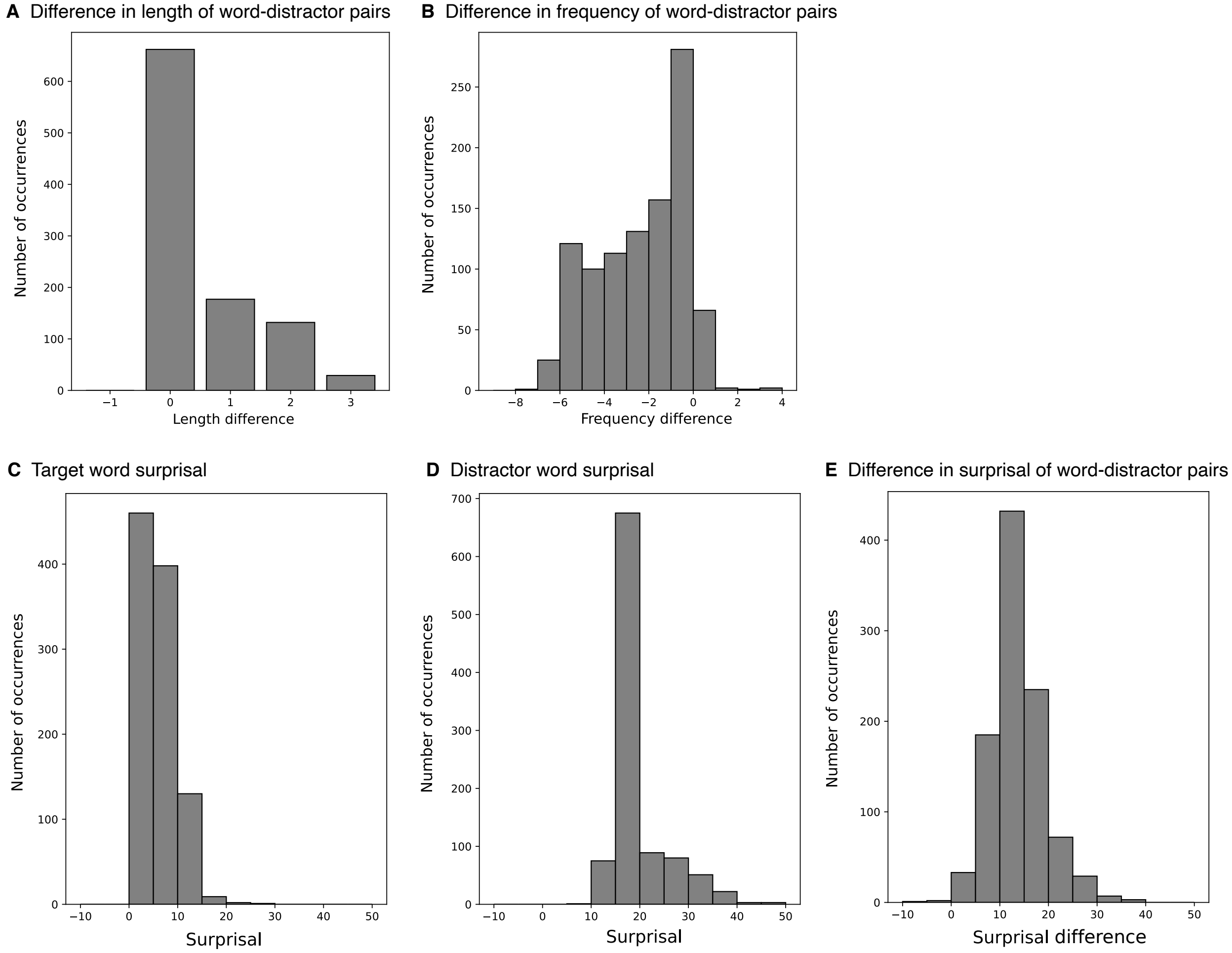

We quantified the quality of the automatically generated distractor words in the A(uto)-Maze paradigm (see Methods; Word-by-word reading times). The goal was to closely match distractors to target words in terms of word length and lexical frequency, while maximizing the contextual surprisal of the distractor to make it a poor continuation of the sentence. Panel **A** shows the character length difference between the target word and distractor word. Most distractors (66.2%) were matched exactly in length, while some shorter target words were paired with a distractor word that was 1-3 characters longer. Panel **B** shows the log lexical frequency differences between target and distractor pairs: Most words (50.6%) were matched within +/- 2 log units of the target word, while some distractors had lower frequencies than the target word. Panel **C** and **D** show the contextual surprisal (in bits, estimated using GPT2) for the target words and distractor words, respectively. Panel **E** shows the surprisal differences between target and distractor pairs: All distractors except 3 (0.3%) had surprisal estimates that were greater than the target word, and the difference in surprisal between the target word and the distractor was 13.72 bits, on average.

#### SI 14: Behavioral experiment: Concreteness of sentence meanings

This supplement provides additional information about the behavioral experiment targeted at assessing the concreteness of a sentence's meaning.

##### SI 14A: Experimental instructions

In this survey, you will be asked to rate 100 sentences. We would like you to rate each sentence according to **how concrete the sentence's meaning is** on a scale from 1 (not at all) to 7 (very much). Concrete meanings are tangible and can be experienced through the senses—vision, audition, smell, touch, etc.

For example, sentences like *"The sports car's engine roared loudly."* or *"The grandmother peeled a green apple."* describe meanings that are tied to perceptual experience (you can hear the sound of the car's engine, picture the grandmother, etc.), so you might rate them as a 6 or a 7. In contrast, sentences like *"Honesty builds strong bonds."* or *"I carry regret for missed chances."* express meanings that are not directly tied to perceptual experience, so you might rate them as a 1 or a 2.

<u>Question phrasing for each item:</u>

- How much is the sentence's meaning tied to perceptual experience?

##### SI 14B: Participant exclusion criteria

The following criteria were defined prior to the study (Tuckute et al. 2024). Participants were excluded based on:

1. **Native speaker status**: Participants were excluded based on their native speaker status self-report as well as the Prolific language and location filters.
2. **Sentence completions**: Participants were excluded if their sentence completions were ungrammatical, contained spelling errors (that were not obvious typos) or if the completions were deeply nonsensical (see SI 14C).
3. **Average response time**: Participants were excluded if the average response time per question was less than 3 seconds.
4. **Lack of variance in ratings**: Participants were excluded if they only used a total of 2 unique rating values (out of 7) for all items in the survey.
5. **Correlation with other participants**: Participants were excluded if the average Pearson correlation with the ratings of remaining participants fell below 2 standard deviations below the mean inter-participant correlation. The inter-participant correlation was computed by correlating a vector of responses for a given participant with the vector of responses for each of the remaining participants and taking the average of these pairwise correlation values.

##### SI 14C: Sentence completion prompts

Participants were instructed: "Please finish the following sentences:".

1. The garden that …
2. When I was younger, I would often …
3. The longer the workers protested, ...
4. I could never have imagined that …
5. Because Jane lived by herself, …
6. The most difficult thing about the trip was that ...
